# How bending scales: Geometric scaling differentially affects force responses in bending biomechanical structures

**DOI:** 10.64898/2026.10.19.760265

**Authors:** Michael J. Bennington, Stephen M. Rogers, Gregory P. Sutton, Hillel J. Chiel, Victoria A. Webster-Wood

## Abstract

The size of an animal’s body greatly affects which biomechanical forces (muscle forces, elasticity, viscosity, and inertia) dominate its behavior. By knowing how the relative contributions of these forces scale with the animal’s size and geometry, we can make predictions about the neural control required to generate its behaviors and to respond to perturbations. The scaling laws of force-producing and energy-transforming structures utilized in neuromechanics focus on spring-like muscles and elastic structures – those that generate forces by changing their length. However, many systems generate and transform energy by bending their tissues, that is, changing their curvature. Even though bending structures are common in biomechanical systems, their scaling laws have not been thoroughly studied. In this paper, we investigate how the forces produced by bending structures scale with the size and geometry of an animal and how that scaling differs from standard spring-like muscles and elastic elements. Through analytical and computational investigations, we demonstrate that particular allometries yield different force scaling laws for bending structures. These different scaling laws impact the duration of quasistatic behaviors, may place limits on animal size, and provide material-efficient methods of remodeling in energy-transforming systems. Together, this work demonstrates the differences between bending and straining biomechanical structures and the need for integrative approaches to scaling analysis that account for specific morphologies and deformations, not just a structure’s cross-sectional area.

## 1 Introduction

Bending structures and movements underpin behaviors in many animals (Fig. 1). For example, the body may be thrown into a series of waves as demonstrated by nematodes, eels, and snakes [37]. Alternatively, in locomotion, leeches, maggots, and geometrid caterpillars perform an inching motion by flexing and straightening the entire body [29]. Particular body parts can be muscular hydrostats that move, at least in part, by bending without fixed joints [22]. These structures include the arms of cephalopods, the trunks of elephants, and vertebrate tongues. In some extremely fast behaviors, elastic energy is stored through the bending of specialized cuticular structures prior to movement [25]: for example, those associated with latch-mediated jumping in insects such as the semilunar process of grasshoppers [30] or the pleural arch of planthoppers [4, 36]; or powering the rapid strikes of trap jaw ants [42], mantis shrimps [33], and pistol shrimps [26].

**Fig. 1:**
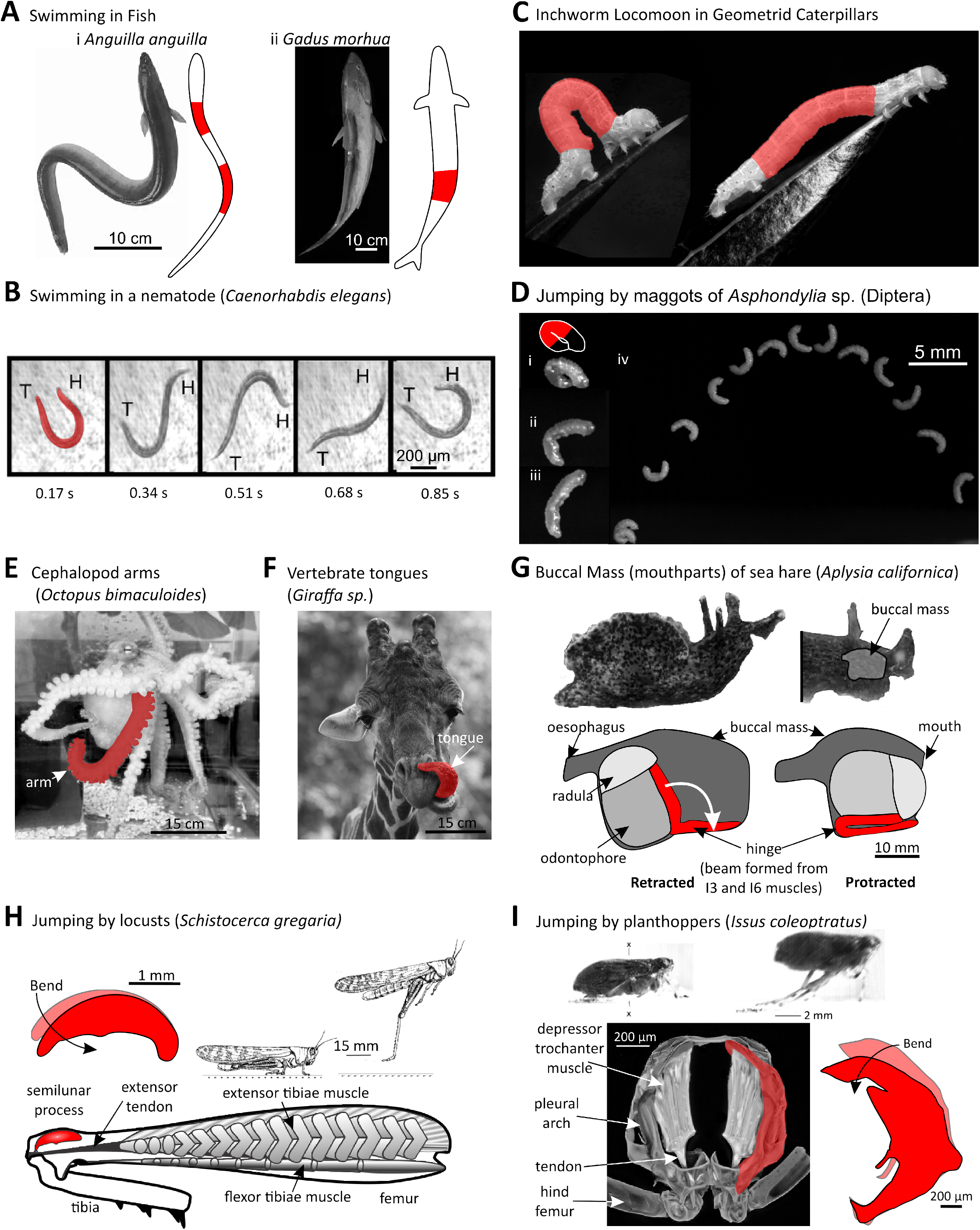
Bending beam-like structures underpin many behaviors. Body regions that can act like bending beams are shown in red. A) Different degrees of body bending underlying different modes of swimming in fish, for example, the i) European eel and ii) Atlantic cod (derived from Di Santo et al. 2021 [35]). B) At a much smaller scale, swimming in the nematode *C. elegans* also involves extensive bending of the body (H indicates the head and T the tail; data from Ghosh and Emmons 2006 [13]). C) The ‘inchworm’ locomotion characteristic of Geometrid caterpillars and leeches requires that the body be bent into a loop [29] (Source images by Katja Shulz, Creative Commons 2.0 license). D) Some Dipterous maggots, including the fungus gnat (Cecidomyiidae) *Asphondylia* can jump by first bending and locking their bodies into a closed loop (data from Farley et al. 2019 [9]). E) The highly flexible arms of cephalopods perform many functions, including locomotion and feeding (picture from Wang and Ragsdale, 2018 [44]). F) The tongues of vertebrates are flexible muscular hydrostats with bending beam-like properties [3] (Source image by Till Niermann, Creative Commons 3.0 license). G) The mouthparts (buccal mass) of the sea slug *Aplysia californica* can perform a number of different feeding behaviors in which a grasper (odontophore) is projected through its anterior lumen towards (and away from) the mouth. Several structures within this muscular hydrostat, including the hinge, behave like bending beams (modified from Bennington et al. 2026 [3]). H) The more familiar jump of the locust also involves the storage of elastic energy by bending the high-stiffness semilunar process located at the distal end of the hind femur. The recoil release of the semilunar process powers high-velocity jumps (modified from Rogers et al. 2016 [32]; Burrows et al. 2021 [4]). I) Likewise, jumping by the true bugs within the Auchenorrhyncha involves the prior bending of the pleural arch, a cuticular structure within the body wall, to store elastic energy (modified from Siwanowicz and Burrows, 2017 [36]; Burrows et al. 2021 [4]). Images have been gray-scaled, and backgrounds have been adjusted to highlight key subjects.

Bending, however, has often been neglected in the scaling analysis of behaviors except in discussions related to constraints on animal (or plant) size imposed by limits on yield stress, buckling instability, or similar indicators of incipient structural failure [28, 27]. While these analyses are fundamental, they do not directly address the role of structures that are intentionally bent to transform energy. Instead, they focus on how the different dimensions of a structure should scale relative to each other to maintain structural integrity. Additionally, the analysis of beams is typically restricted to scaling laws that are isotropic with respect to the beam’s cross-section, neglecting how bending direction could affect scaling [20]. Do the forces from bending-dominated structures scale in the same way as linear tension-springs? If not, how does anisotropy in the geometric scaling law in bending structures govern force production and ultimately impact the behaviors of the animals that utilize them?

In this article, we investigate how the forces produced by bending-dominated biomechanical structures scale with a system’s mass and what impact those forces have on behavior. Based on an analysis of the energetics of beams, we derive a general scaling law for the force produced by structures that bend. This relationship reveals that under particular allometries related both to directions of the cross-section and the length of the structure, bending and straining structures do not produce the same force scaling law even when they share the same geometric scaling law. We validate the qualitative predictions of the scaling analysis using computational experiments with a simplified beam model under various geometric scaling laws. Next, we generalize these findings to the analysis of a simple quasistatic system, in which the passive elastic forces of tissues are the dominant forces that muscles must oppose, and thus any deviations from predicted scaling laws are most likely to impact how behaviors are generated. We use this simplified model to investigate how beam-like force scaling affects the durations of behaviors. Finally, we discuss the implications of non-spring-like force scaling on the behaviors, growth, and remodeling of biological systems. Together, this work broadens the scope of possible scaling force relationships to expect in biological systems and lays the foundation for future mechanical, anatomical, and behavioral scaling investigations of bending biomechanical structures.

## 2 Methods and Model Derivations

### 2.1 Variable convention

Throughout this article, italicized letters (Latin or Greek) denote scalar values. Vectors are stylized with an arrow 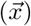, and unit vectors with a hat 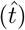. The expression 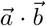 denotes the vector dot product between vectors 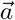 and 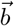, which produces a scalar quantity.

### 2.2 Overview of bending mechanical structures

Before we get too far into the mathematical analysis of how bending elements may influence biomechanical systems, it is helpful to establish some intuition about beams. For this, let us consider an object many of us have bent in our lives: the common ruler (Fig. 2(a)). We can align three perpendicular axes with the ruler: the long edge of the ruler aligns with the axial axis (red; labeled with the coordinate *s*), the broad flat width of the ruler is measured along the *α* axis (teal), and the thin narrow edge is measured along the *β* axis (light blue). When we hold both ends of the ruler, we find that it is much easier to bend around the *α* axis (Fig. 2(a2)), where we are bending perpendicular to the thin edge. If instead we rotate the ruler and try to bend around the *β* axis (Fig. 2(a3)), where we are trying to bend against the full width of the ruler, it becomes much more difficult. In either direction, the amount of torque (or moment) we need to apply to the ends of the ruler to bend it further increases as the ruler becomes more and more curved. This simple intuition tells us that the torque (or moment) needed to bend this beam-like object is going to be somehow proportional both to the thickness of the object perpendicular to the bending axis and to the curvature of the bend. Let’s now formulate this more mathematically to establish our model.

**Fig. 2:**
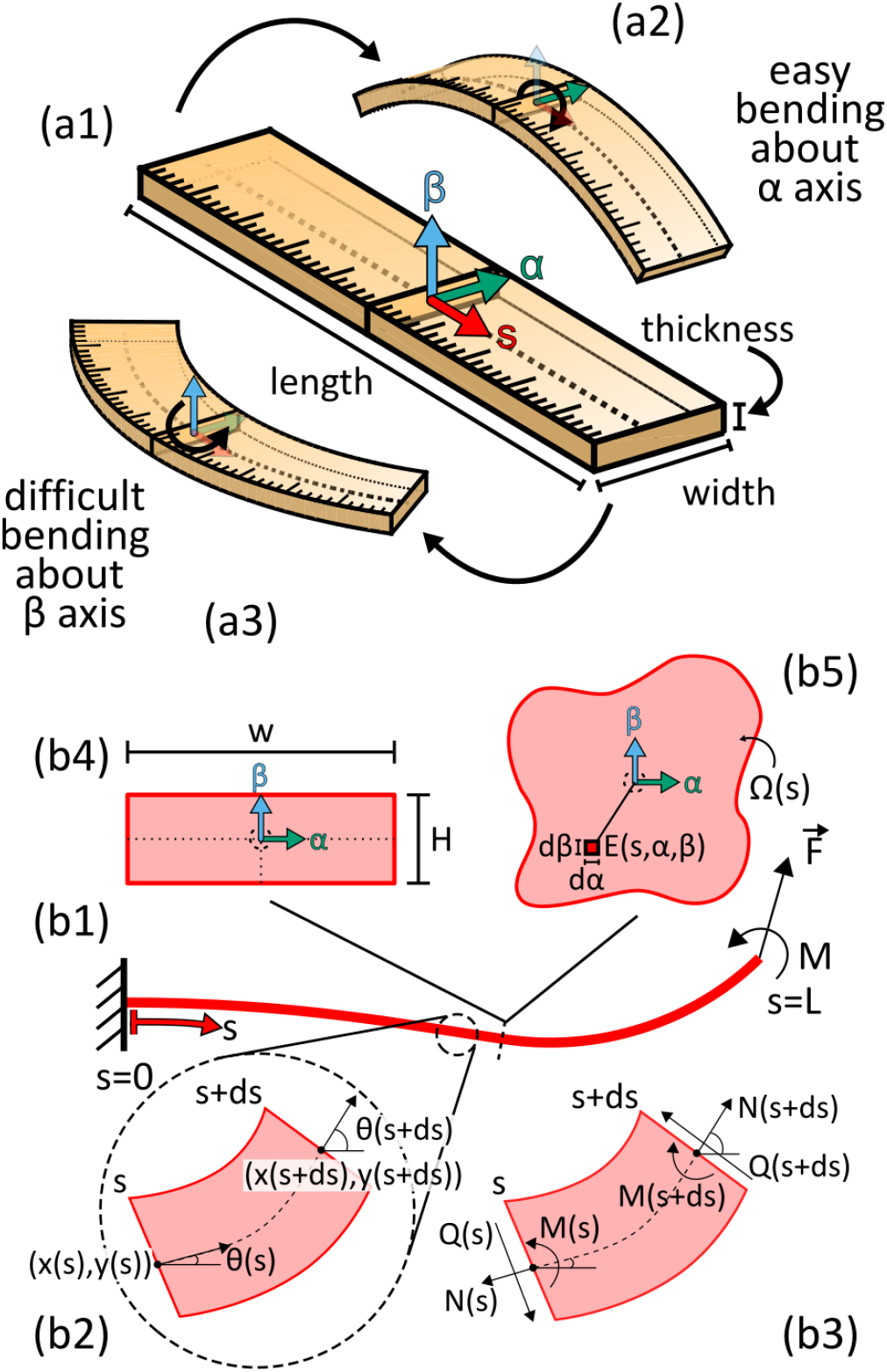
Primer on bending mechanical structures and multi-variable integration. (a1) A common example of a beam-like structure is a ruler. (a2) Bending the ruler about the axis parallel to the width of the ruler (α) is easier than (a3) bending about the axis parallel to the thin edge (β). This reflects that, not just the cross-sectional area (CSA), but how far from the bending axis the material of the cross-section lies impacts the force required to bend a beam. (b) Mechanics of a general beam structure. (b1) A beam is an elastic structure defined by a 1-dimensional curve that resists both stretching and bending. (b2) The position of a beam (x(s), y(s)) and its heading angle θ(s) is defined at each position (s) along the length of the beam. (b3) The beam’s position under an applied load is determined using a force/moment balance on each infinitesimal section of the beam. N(s): axial force, Q(s): shear force, M(s): bending moment. (b4-b5) The axial and bending stiffness of a beam are determined by the material stiffness (Young’s modulus, E) and the cross-sectional geometry. (b4) For the main analysis presented here, we assume the beams have a rectangular cross section with width w and thickness H. (b5) The same scaling laws can be derived for beams of arbitrary cross-sectional geometry (Ω(s)) and stiffness distribution (E(s, α, β)).

### 2.3 Mechanics and scaling of elastic biomechanical structure

In this article, we will compare the scaling of two different types of biomechanical structures. The first, which we call “spring-like,” only resists stretching (changes in length), and the force generated by it acts along its length. That is, they act like tension springs. The second, which we call “beam-like” or “beams,” resists both stretching and bending (changes in curvature), and both forces and moments (or torques) are generated passively by deforming the structure (Fig. 2(b)). These are analogous to the ruler described above. Thus, a spring-like structure can be seen as a beam where the bending stiffness can be neglected. Both of these structures are defined by a (topologically) 1-dimensional curve passing through the middle of their 3-dimensional geometry. Here, we only consider planar structures with linear constitutive equations for simplicity, but a similar analysis could be applied to non-planar deformations and nonlinear constitutive equations. For these planar deformations, all bending will occur about the *α* axis.

Briefly, the quasistatic governing equations for each structure are derived by applying a force balance (and moment balance, for beams) on each infinitesimal segment along the line defining the structure (Fig. 2(b3), see Supplemental Materials S2) [2]. These force (moment) balance equations are then coupled with constitutive equations defining the relationship between the forces (moments) that these structures apply and the deformation that they undergo. For springs and beams with linear constitutive relationships, the force along the length of the structure is

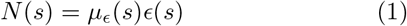

where *s* is the arc length position along the structure, *N*(*s*) is the axial force, *µ*_*ϵ*_(*s*) is the axial stiffness (units of force), and 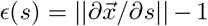 is the axial strain. For beams, the bending moment is

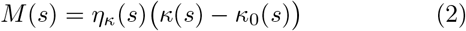

where *M*(*s*) is the bending moment, *η*_*κ*_(*s*) is the bending stiffness (units of force *×* length^2^), and *κ*(*s*) = ∂*θ/*∂*s* is the beam’s curvature. *κ*_0_(*s*) is the curvature in the rest position.

Both the axial and bending stiffnesses depend on the structure’s cross-sectional geometry and Young’s modulus, *E* (Fig. 2(b4)). For the simple case where the Young’s modulus is constant throughout the cross section, axial stiffness (*µ*_*ϵ*_) is equal to the cross-sectional area (CSA) times the Young’s modulus (*µ*_*ϵ*_ = *EA*), and the bending stiffness (for bending about the *α* axis) is equal to the Young’s modulus times the second polar moment of area 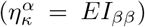 [2]. This second polar moment of area is analogous to the variance of a dataset and captures how spread out (dispersed) the cross-section is in the direction perpendicular to the bending axis. This also explains why, in our example of the ruler, bending is easier about the *α* axis, because the cross section is less spread out from the *α* axis than it is from the *β* axis (Fig. 2(b4)). Intuitively, one can relate the axial stiffness to the total amount of material in the cross section and the bending stiffness to how spread out that material is from the axis of rotation.

For this analysis, we will look at the specific example of a rectangular cross-section. However, the same analysis for an arbitrary cross section is found in Box 1. For a rectangular cross section with width *w* in the *α* direction and thickness *H* in the *β* direction (Fig. 2(b4)), the axial stiffness is

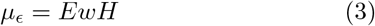

and the bending stiffness is

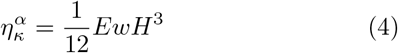

For the bending stiffness about the *β* axis, *w* and *H* exchange their exponents.

As the geometry of these structures scales, the axial and bending stiffnesses also change. Here, we assume the lengths of these structures scale as a power law of a characteristic mass (*m*). This could be the mass of the whole animal or of the component that contains the elastic structure of interest. Let 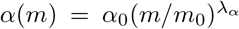 and 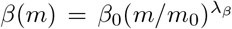, where (*α*_0_, *β*_0_) are the coordinates in the reference system with mass *m*_0_. Here, we assume that the Young’s modulus distribution does not change with mass, implying no meaningful structural remodeling or morphological changes occur during growth. For a brief example of scaling under a morphological change, see the Appendix. Using these scaling laws, we can determine how stiffness scales with the mass of a structure:

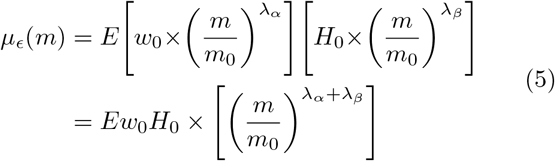

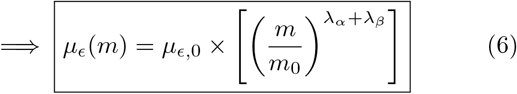

and

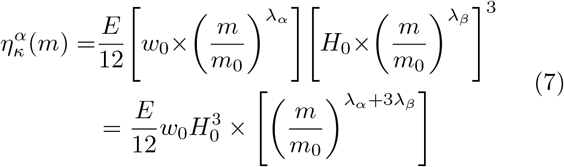

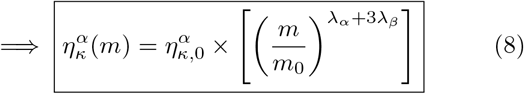

The bending stiffness for bending about the *β* axis will have the same form as 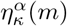, but the scaling parameter will be 3*λ*_*α*_ +*λ*_*β*_. Here, *µ*_*ϵ*,0_ and 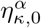 are the stiffness values in the reference system where *m* = *m*_0_.

Under isometric scaling (*λ*_*α*_ = *λ*_*β*_ = 1*/*3), axial stiffness scales as mass^2*/*3^. This aligns with traditional scaling analyses of mechanical structures in which axial forces scale identically to the CSA (*A ∝* 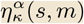). However, bending stiffness scales as mass^4*/*3^. Through our theoretical and computational investigations, we determine the implications of this supra-force scaling of bending stiffness in beam structures. These scaling relationships also show that, while the axial stiffness is isotropic (i.e., the *α* and *β* directions contribute equally), the bending stiffness is anisotropic. The direction perpendicular to the axis of rotation (*β*) is more heavily weighted than the direction parallel to it (*α*), contributing to stiffness as 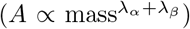 as compared to 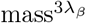, respectively. The implications of this anisotropy, and its interactions with allometric scaling laws, will also be investigated.

#### Box 1

**Stiffness parameter scaling law for arbitrary crosssectional geometries**

In the main text, we derive parameter scaling laws for a beam with a rectangular cross-section as an illustrative example. The same scaling laws can be derived for an arbitrary cross-sectional geometry. For a structure with an arbitrary cross-section Ω(*s*) (Fig. 2(b5)), let the local coordinates (*α, β*) have their origin at the centroid of the cross-section. *E*(*s, α, β*) is the Young’s modulus distribution on the cross-section. Axial stiffness is found by integrating the Young’s modulus across the cross-section [2]

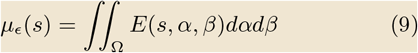

For a primer on the multi-variable integration, see Supplemental Materials S1. This is equivalent to the average Young’s modulus times the CSA. The bending stiffness for bending about the *α* axis is found as the second moment of the stiffness distribution [2]

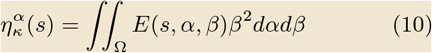

The analogous bending stiffness for bending about the *β* axis takes the same form with the *β*^2^ switched to *α*^2^. For simple geometries (e.g., rectangles, circles, annuli) and a uniform Young’s modulus, these expressions reduce to the Young’s modulus times area (for axial stiffness) and Young’s modulus times the second polar moment of area (for bending stiffness).

Under the same geometric scaling law assumptions from the main text (no structural remodeling and lengths scale as power laws of a characteristic mass) in the expressions for the axial and bending stiffness (Eqns. 9 and 10, respectively), the scaling relationships for these stiffnesses become:

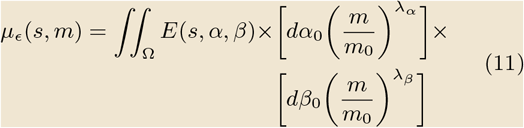

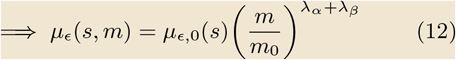

and

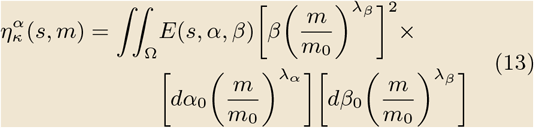

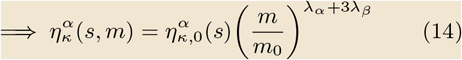

The bending stiffness for bending about the *β* axis will have the same form as 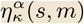, but the scaling parameter will be 3*λ*_*α*_ + *λ*_*β*_. **Note:** these scaling relationships are identical to the ones derived for the rectangular case; the rest configuration stiffnessess (*µ*_*ϵ*,0_, *η*_*κ*,0_) will change, but the scaling laws are cross-section geometry agnostic.

### 2.4 Force scaling analysis of a bending-dominated structure

To investigate how different geometric scaling laws impact the forces produced by bending-dominated structures, we developed a simplified model of a passive bending structure modeled as a beam (for details, see Supplemental Materials S3). The beam’s rest geometry, while not directly based on any particular biological structure, was defined by a cubic Bézier spline to approximate structures seen in animals, like the pleural arch of plant hoppers and the semilunar process in grasshoppers (Fig. 3(a,b)). Importantly, this particular model was used for a demonstrative investigation, but the general conclusions of this analysis are geometry-agnostic. The beam is fixed at one end, and a force (approximating a muscle force on the structure) is applied to the other (Fig. 3(c1)). The beam is constrained to the plane, such that it can displace in the *x* and *y* directions and rotate about the beam’s *α* axis. For simplicity, we assume the beam has a rectangular cross-section with the same aspect ratio along its full length (Fig. 3(c2)). In simulation, we can measure the relationship between the muscle’s length (which sets the beam’s displacement) and the reaction force produced by the beam to produce force-displacement curves.

**Fig. 3:**
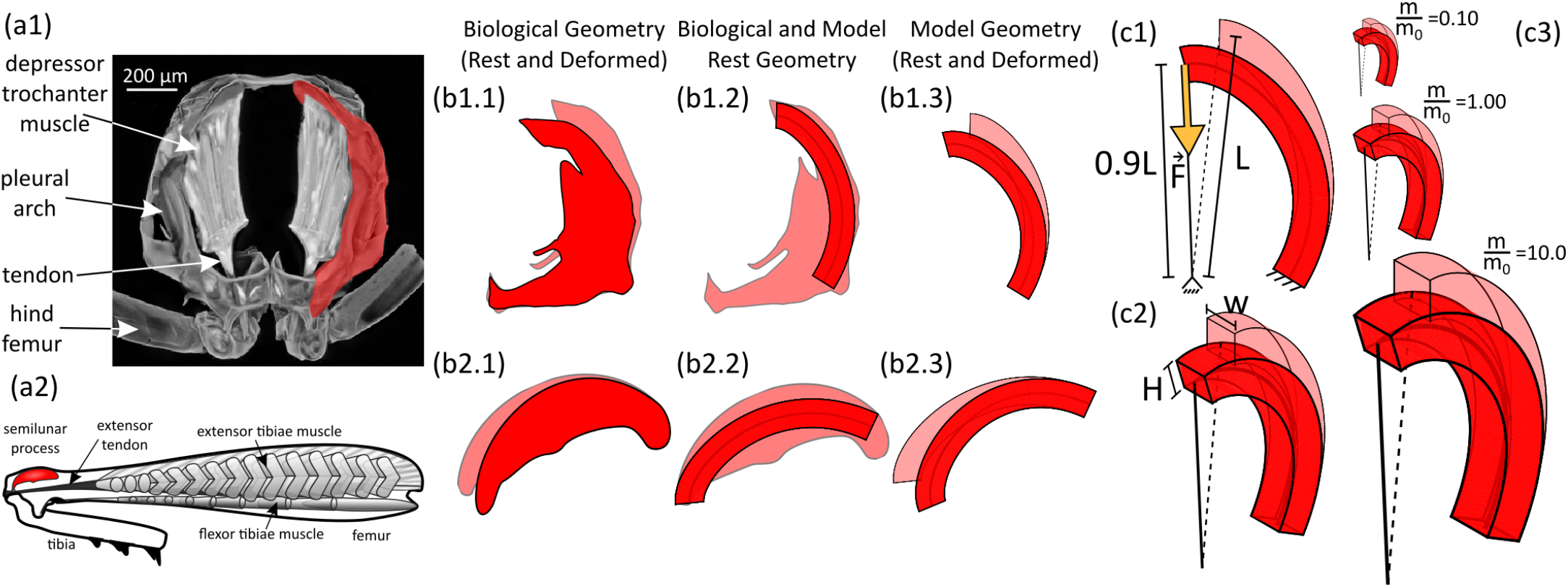
General model of a bending-dominated structure. To investigate the effects of geometric scaling on a beam-like biomechanical structure (like the (a1) pleural arch of a plant hopper or (a2) semilunar process of grasshoppers), we developed a general model of a bending structure using beam mechanics. (b) The beam’s geometry is inspired by the pleural arch and semilunar process, but the conclusions based on the simulations are system-agnostic. Additionally, the specific geometry and loading conditions do not fully match those of the pleural arch or the semilunar process. Results should not be interpreted as simulations of these structures. The same beam geometry (rescaled and rotated to match the biological morphology) is used in (b1) and (b2). (c) The simulated beam is loaded at its tip by a force (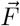, (c1), pointing towards a fixed point near the beam’s base. This is analogous to a muscle or tendon force acting to bend this structure. The beam’s geometry is determined by (c1) the rest length of the “muscle” applying the force to the beam (*L*) and (c2) the width (*w*) and thickness (*H*) of the rectangular cross section. (c3) Rescaled models were created for mass ratios (*m/m*_0_) ranging from 0.1 to 10. Models in (c3) are shown to scale. Note: the images in (a1) and (a2) are reproduced from Fig. 1. See that caption for details of these images.

The beam’s geometry is characterized by three parameters (Fig. 3(c1-c2)): the resting length of the “muscle” (*L*), the width of the cross section parallel to the bending axis (*w*), and the thickness of the cross section perpendicular to the bending axis (*H*). *L* fully defines the beam’s centerline geometry (See Supplemental Materials S3). To study the effects of geometric scaling, we created rescaled versions of this model, where these three parameters were scaled as power laws of the model’s mass relative to a reference mass (e.g.,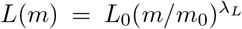). These rescaled anatomies were created for mass ratios (*m/m*_0_) ranging from 0.1 to 10 (Fig. 3(c3)). The scaling parameters for the three directions took on values in the range *λ*_*α*_, *λ*_*β*_, *λ*_*s*_ *∈* [0.25, 0.5]. This range was chosen to cover both sides of isometry and to span scaling values observed in the literature. Force-displacement curves for each rescaled model were then simulated and compared to determine the force scaling relationship under different geometric scaling laws.

### 2.5 Scaling of a quasistatic behavior

To investigate the behavioral impact of beam-like structure scaling, we constructed and analyzed a simplified model of a quasistatic agonist-antagonist system. We chose to investigate a quasistatic system because the dominant forces constraining its behavior are elastic forces. Thus, any scaling differences in the beam-like elastic structures will impact the scaling of behavior. This model is intentionally animal-agnostic, as its qualitative predictions apply to many quasistatic systems. Additionally, the model was formulated with one translational degree of freedom (1 DoF), but the same analyses could be conducted on a 1 DoF rotational model, e.g., a leg joint, and the qualitative prediction are compatible (See Appendix 6.1).

For this model, we assume a mass that can translate along a single axis (Fig. 4). Let the normalized position of this mass be *q*∈ [0, 1]. This mass is acted upon by an agonist/antagonist muscle pair, labeled muscles 1 and 2. Both muscles produce passive and active forces. The passive forces encompass those generated by the muscle itself and any other passive structures acting in parallel. Let the passive force generated by muscle *j* ∈ [1, 2] be 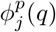, and its active force be 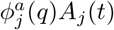, where *A*_*j*_(*t*) *∈* [0, 1] is the normalized activation of that muscle. Here, 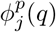 and 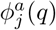 capture the overall axial force produced by muscle *j* at a given position, incorporating length dependencies and position-dependent mechanical advantage terms. In this model, we refer to forward motion (increasing *q*) caused by the agonist as “protraction,” and backwards motion (decreasing *q*) caused by the antagonist as “retraction.”

**Fig. 4:**
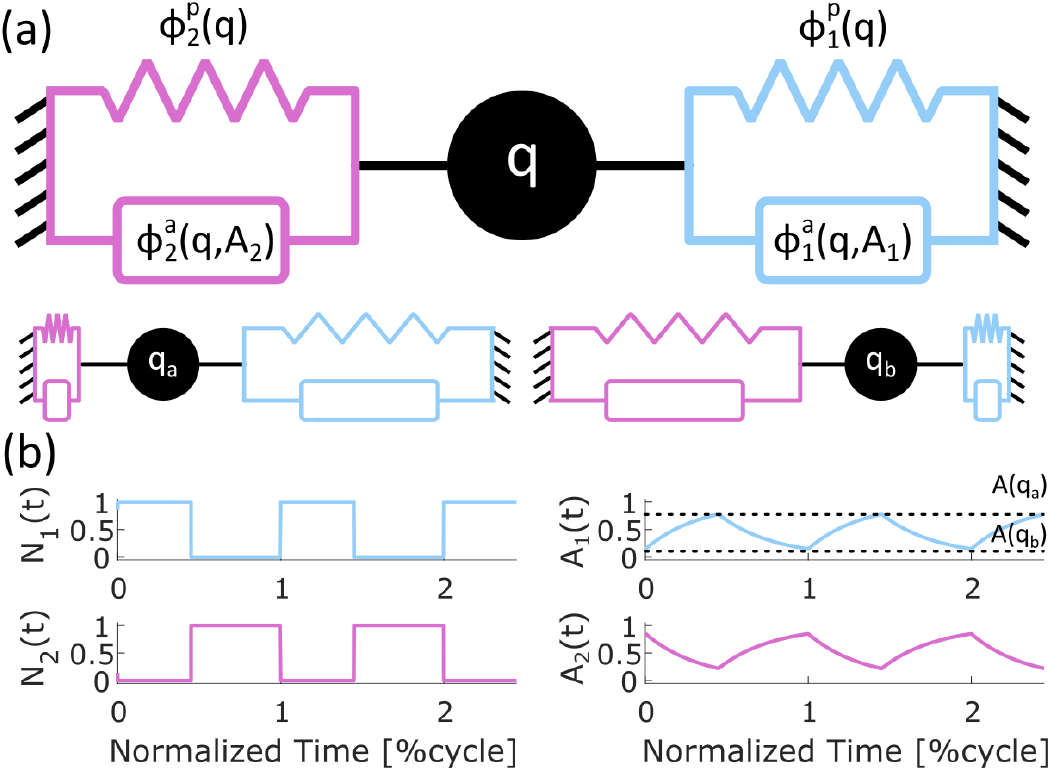
Model of a uniaxial 1-degree-of-freedom (1DoF) quasistatic system. (a) To investigate the effects of beam-like force scaling on the behavior of quasistatic systems, we developed a simplified 1DoF quasistatic translational model. Here, an agonist (blue) and antagonist (pink) muscle pair act on a body that can translate between normalized positions *q*_*a*_ and *q*_*b*_ (bottom) during steady-state behaviors. We set up the general framework for this agonist-antagonist muscle system, and below we will investigate how replacing a spring link muscle with a bending beam-like muscle affects scaling. The muscles generate both passive 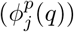 and active 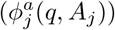 forces to move the body (*j* ∈ [1, 2]). (b) We assume that the muscles receive input stimuli (*N*_1_ = 1 *− N*_2_) out of phase from each other. These neural inputs are then passed through low-pass-filter-like muscle activation dynamics to produce the activations *A*_1_(*t*) and *A*_2_(*t*) = 1 *− A*_1_(*t*).

At each time point, the agonist and antagonist forces are balanced:

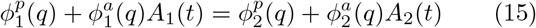

Because the model is quasistatic, any inertial or viscous forces are negligible and, therefore, omitted. Next, we assume muscle activation (*A*_*j*_(*t*)) acts as a first-order filter [45] of a normalized neural input *N*_*j*_(*t*) *∈* [0, 1]. Thus, activation is governed by the differential equation:

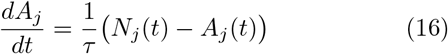

where *τ* is the time constant of the muscle activation dynamics. Finally, we assume the muscles receive neural inputs out of phase with each other (i.e., *N*_1_(*t*) = 1 *− N*_2_(*t*)). Therefore, in steady-state cycles,

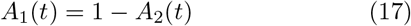

(Fig. 4(b); see Supplemental Materials S5). Plugging Eqn. 17 into Eqn. 15, we can solve for the agonist activation as a function of position:

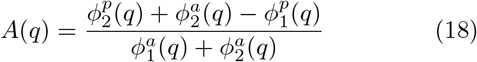

Here, we drop the subscript “1” for simplicity. This is the level of agonist activation required to achieve the position *q* during a steady-state cycle. The muscle activation at position *q* is fully determined by the forces at that position under steady-state conditions. But the time course of activation (and, therefore, behavior) is determined by the activation dynamics (Eqn. 16).

During steady-state behaviors, the system oscillates between *q*_*a*_ and *q*_*b*_ (Fig. 4(a, bottom)), representing the minimum and maximum positions reached during the behavior. By Eqn. 18, this implies the activation will oscillate between *A*_*a*_ = *A*(*q*_*a*_) and *A*_*b*_ = *A*(*q*_*b*_) (Fig. 4(b)). If we assume that, during protraction (*q*_*a*_ *→ q*_*b*_), the neural input to the protractor muscles is maximum (*N*_1_ = 1) and that during retraction (*q*_*b*_ *→ q*_*a*_), the neural input is minimal (*N*_1_ = 0), then the duration of these phases are solved analytically as functions of *A*_*a*_ and *A*_*b*_ (see Supplemental Materials S6). This yields that the duration of protraction (*T*_*p*_) is

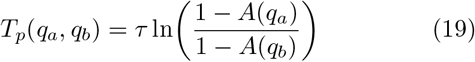

and the duration of retraction (*T*_*r*_) is

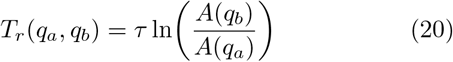

The total cycle duration (*T*_*c*_) is then *T*_*c*_ = *T*_*p*_ + *T*_*r*_. As the neural inputs were always at their maximum value during their respective phases, these expressions serve as lower limits on the segment durations, but similar expressions could be derived for a different known neural input signal. These cycle times are fully determined by the extreme positions of the cycle, the force response of the constituent muscles, and the time constant of the muscle activation. Thus, if animals of different sizes share the same normalized extreme positions, the scaling of behavior durations can be fully determined by the scaling of the muscle activation time constants and of the muscle forces. Using Eqns. 18, 19, and 20, we will investigate the effects of spring- and beam-like scaling on the duration of behaviors.

## 3 Results

### 3.1 Isometric scaling of beam structures leads to spring-like scaling

How does the supra-force scaling of a beam’s bending stiffness impact the force-displacement relationship of a bending-dominated biomechanical structure? Using a simple beam model as a test case (Fig. 3), we first investigated the impacts of isometric growth in a beam’s cross-section. For this, we scaled the model’s axial and bending stiffnesses by mass^2*/*3^ and mass^4*/*3^, respectively (Fig. 5(a); see “Mechanics and scaling of elastic biomechanical structure“). Under these conditions, the force at any normalized displacement level increased monotonically as a function of mass (Fig. 5(b)). However, when we normalize the forces by mass^2*/*3^ (the CSA scaling rate and the expected scaling rate for an isometrically-scaled spring-like structure), all force curves collapse to the same normalized one (Fig. 5(c)). Additionally, when we observed the force level at two displacements (5%L and 10%L) at each mass level, we see the force scales exactly as mass^2*/*3^ (Fig. 5(d)). Thus, under isometry, despite bending stiffness scaling faster than axial stiffness and bending being the dominant energetic component in these beams, the force-displacement relationship still scales identically to a spring-like structure. How do these components with different scaling relationships in their stiffness lead to the same scaling relationship in their emergent force behaviors?

**Fig. 5:**
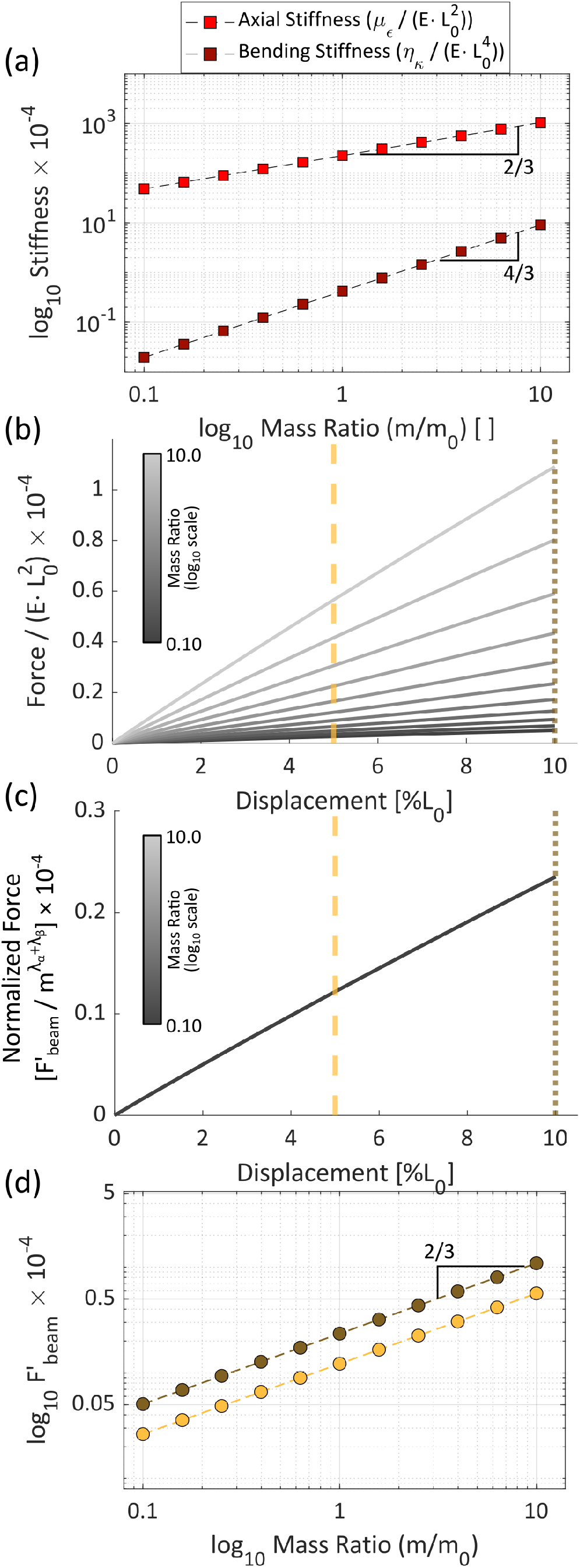
Isometric scaling of a bending-dominated structure. (a) Axial (*µ*_*ϵ*_) and bending (*η*_*κ*_) stiffness of the modeled beam under isometric growth. The slopes indicated in the log-log plot correspond to the power-law exponent for the stiffness scaling laws. Both stiffness values are de-dimensionalized using the reference configuration geometry (see legend) so that both are unitless. (b) In simulated force-displacement experiments, the beam force at any normalized position increases monotonically with mass. (c) When the forces are normalized by the expected scaling law of a spring-like structure (force *∝* mass^2*/*3^), the force curves collapse down to a single curve, indicating the different beam force curves scale as mass^2*/*3^ under isometric cross-sectional scaling. Here, 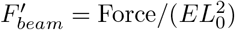. (d) This scaling can also be seen by observing the force at two displacements levels (light yellow: 5% muscle strain; dark yellow: 10%; levels indicated in (b) and (c) by vertical lines of the same color), which both scale in log-log space with a slope of 2/3.

### 3.2 A generalized force scaling relationship for beam structures

To investigate the cause of this observed spring-like scaling for beams, we derived a general scaling relationship for the forces produced by a beam using an energy analysis. For a quasistatic elastic beam fixed at one end with a force 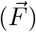 applied at the other end (Fig. 3(c1)), the work done by the force 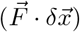 equals the change in elastic energy (*δU*_*beam*_) stored by the structure. This elastic energy can be stored either by stretching (*U*_*s*_) or by bending (*U*_*b*_). Thus,

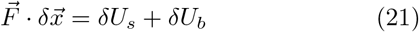

The elastic energy stored by stretching and bending, respectively, are

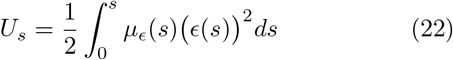

and

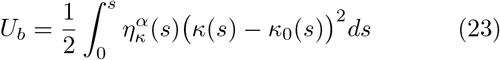

for a given configuration of the beam. These are analogous to the elastic energy of a Hookean spring (*U*_*spring*_ = (1*/*2)*k*(Δ*x*)^2^). Here, *ϵ*(*s*) and *κ*(*s*) are the beam’s strain and curvature distributions under static equilibrium with the force 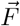. By the Chain Rule, we can expand Eqn. 21 as

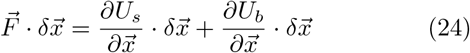

We define 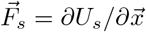 as the force produced by stretching the beam (the “stretching force”), and 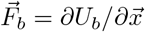 as the force produced by bending the beam (the “bending force”).

With these general expressions for the stretching and bending forces, we can determine a general scaling relationship for the force magnitudes using the scaling rates of the expressions’ components. We drop the vector notation for the forces, as we are only concerned with their magnitudes. The stiffness parameters’ scaling relationships are determined by the geometry of the cross-section (Eqns. 12 and 14). The differential terms (*dx* and *ds*) will both scale like length terms. *dx* scales with the stroke length of the muscle/tissue applying the force to the beam, and *ds* scales as the beam’s arc length. Let *dx ∝* 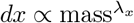 and *ds ∝* 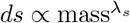. The final terms concern the beam’s deformation. Strain (*ϵ*(*s*) *≈*Δ*L/L*) is the ratio of length change (caused by the stroke length) to the beam’s resting arc length; it therefore scales as 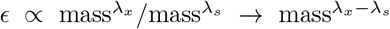. The change in curvature (*κ − κ*_0_ = Δ*κ*(*s*) *≈* Δ*θ/L*) relates to the change in heading angle (caused by the stroke length) distributed over the length of the beam (See Supplemental Materials S4). For small angles, the heading angle change is approximately the stroke length over the arc length (Δ*θ ≈* Δ*x/L*); therefore, curvature (Δ*κ ≈* Δ*x/L*^2^) scales as 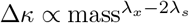.

The constituent terms in the bending and scaling forces expressions combine multiplicatively, so the combined scaling law is found by summing together the scaling parameters of each component. This yields the general scaling relationship for the stretching and bending forces:

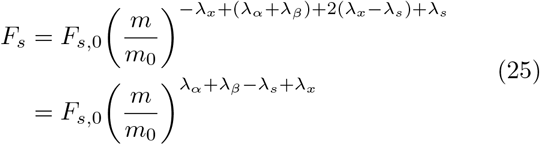

and

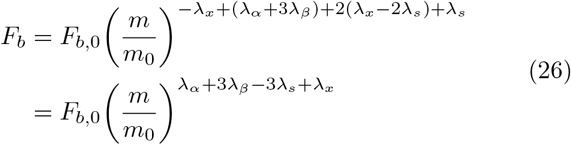

respectively. Note: the term *λ*_*α*_ + *λ*_*β*_ *− λ*_*s*_ in the spring force equations is the scaling of the Hookean spring constant (*k* = *EA/L ∝* 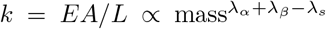 where *F*_*s*_ = *k*Δ*x*). Additionally, the term *λ*_*x*_ *− λ*_*s*_ (the portion not associated with the CSA) accounts for scaling differences in the arc length and stroke length (i.e., if a shorter muscle undergoes a longer stroke length, it will generate more force). If the beam’s arc length and the stroke length grow isometrically (*λ*_*s*_ = *λ*_*x*_ = 1*/*3, and thus, we only consider possible allometry in the cross-section), these expressions reduce to

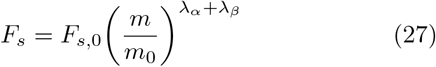

and

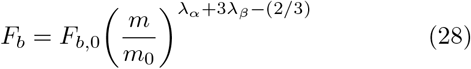

Therefore, the stretching force scales with its stiffness, but the bending force scales more slowly than its stiffness because it depends on the beam’s arc length, both through its effects on curvature and as the moment arm for the applied force. This also implies the stretching force scales like CSA (*A ∝* 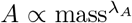 where *λ*_*A*_ = *λ*_*α*_ + *λ*_*β*_) as expected for springs. Finally, the total force from a beam structure (*F*_*beam*_ = *F*_*s*_ + *F*_*b*_) scales not as a simple power law, as predicted from analyses of spring-like structures. Instead, it scales as a weighted sum of power laws:

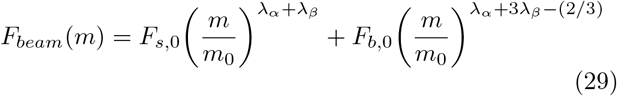

Restricting the cross-section to isometry like in our first simulations (*λ*_*α*_ = *λ*_*β*_ = 1*/*3), we retrieve

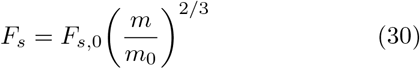

and

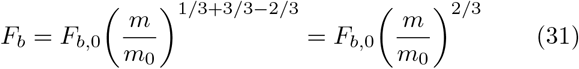

Both forces scale identically, so the total force scales at the same rate:

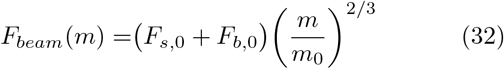

Thus, as observed in the simulation study, under isometry, forces generated by beam structures scale identically to spring-like forces.

### 3.3 Allometric scaling of beam thickness allows for various force scalings

Both stretching and bending forces grow identically to spring-like forces under isometric growth. How do cross-section allometries change these scaling relationships? How does the anisotropy of the bending stiffness scaling law impact these relationships? From the derived scaling relationships (Eqn. 29), we see that, no matter what the scaling law is for the cross-section’s two coordinate directions, the stretching forces will always scale as the CSA, as predicted by standard analyses. The stretching force scaling law also implies that this is true even if the two directions scale differently from each other (*λ*_*α*_ ≠ *λ*_*β*_). This is caused by the symmetry between these two directions in the scaling law; the scaling parameters for these two dimensions contribute equally to the force and CSA scaling laws.

This symmetry does not exist in the bending force scaling law. Instead, the scaling parameter controlling the thickness direction (the direction perpendicular to the bending axis) is more heavily weighted, as it is in the bending stiffness scaling law. The impact of this asymmetry can be seen by factoring out the spring-like scaling term from the total scaling law. If we define the total force at our reference mass as *F*_*beam*,0_ = *F*_*s*,0_ + *F*_*b*,0_, we obtain

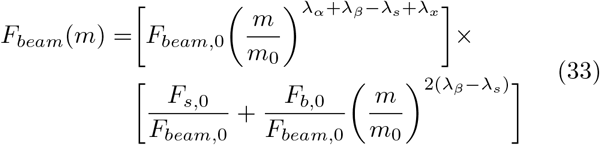

The first bracketed term is the scaling expression for a spring-like structure, where the force scales like CSA (and exactly so when *λ*_*x*_ = *λ*_*s*_). The second bracketed term is a correction term accounting for anisotropy in the bending term and the increased dependence on arc length. As with the bending stiffness term asymmetry, the scaling of the correction term does not depend on the width scaling (*λ*_*α*_). This factored expression allows us to draw four key conclusions about the scaling of bending structures:

1. As the bending force approaches 0 (*F*_*b*,0_ *→*0; i.e., we are dealing with a stretch-dominated, or spring-like, structure), the overall scaling law reduces to that of a spring (*F*_*beam*_ *∝ A*)
2. If the beam’s thickness and arc length scale isometrically (*λ*_*β*_ = *λ*_*s*_ = 1*/*3), or more generally, identically (*λ*_*β*_ = *λ*_*s*_), the bending correction term is exactly 1, and, again, the overall scaling law reduces again to a spring-like scaling law, regardless of *λ*_*α*_
3. By allometrically scaling the beam’s thickness, we get non-spring-like force scaling, with the force scaling faster under faster-than-isometry thickness scaling.
4. Arc length allometry, even if compensated by stroke length allometry (*λ*_*s*_ = *λ*_*x*_ ≠ 1*/*3), will cause non-spring-like force scaling, with the force scaling more slowly under faster-than-isometry arc length scaling.

Thus, the correction term only contributes to bending-dominated systems where the beam’s thickness or arc length scales allometrically.

We tested the latter three of these predictions with a second simulation study (Fig. 6) using the same beam model, in which we ran simulations where one axis was scaled allometrically while the others were fixed at isometry. For all simulations, *λ*_*x*_ = *λ*_*s*_. When the beam’s width scaled allometrically (*λ*_*α*_ *∈* [0.25, 0.5], *λ*_*β*_ = *λ*_*s*_ = 1*/*3), the force from the beam structure scaled identically to a spring-like structure with the same cross-section (Fig. 6(a1)). Thus, when the force was normalized by 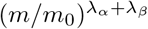 (the expected scaling function for a spring-like structure), all of the scaling curves (for different values of *λ*_*α*_) collapse to the same scale-invariant curve (Fig. 6(a2)).

**Fig. 6:**
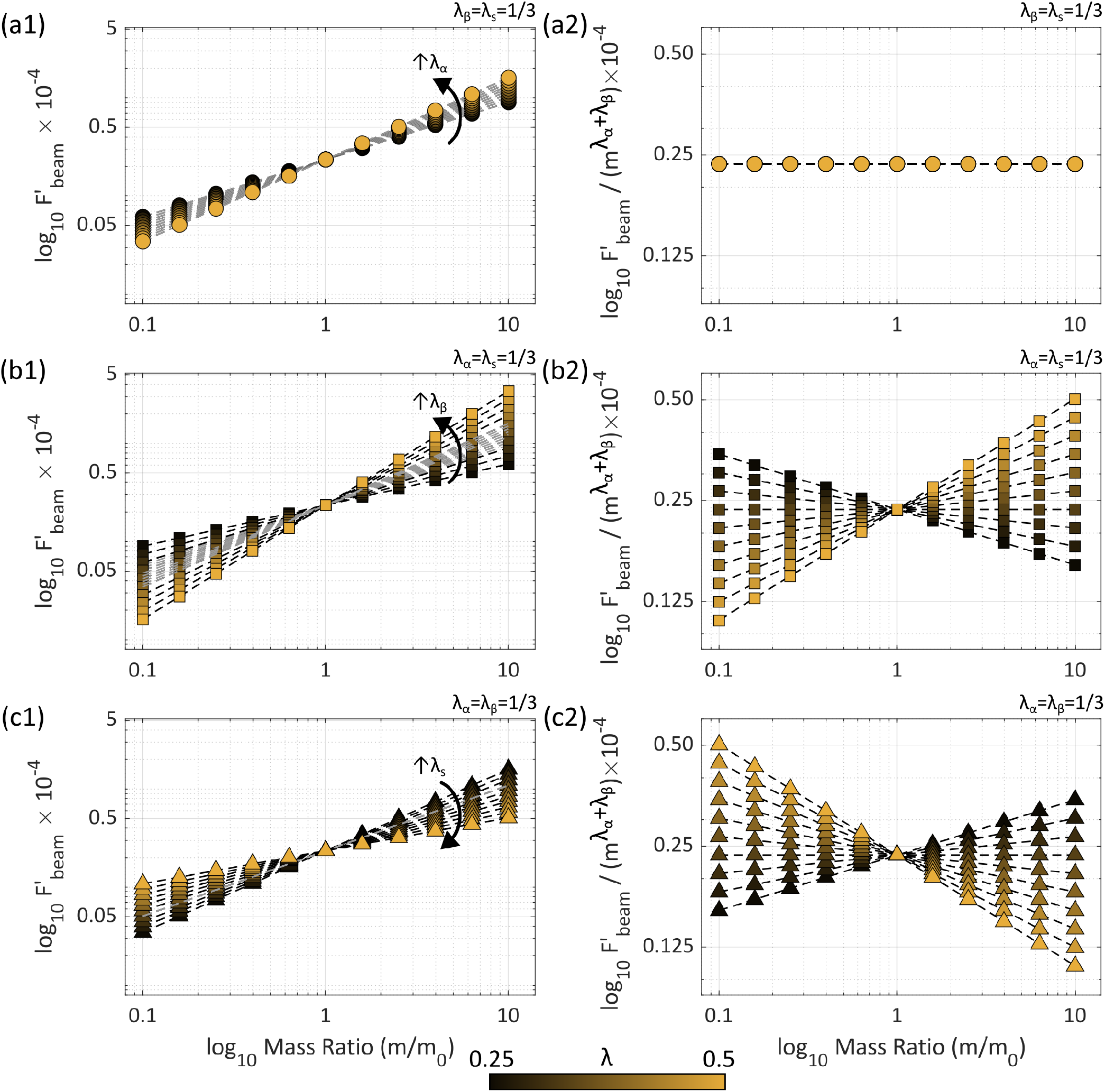
Force scaling of a bending-dominated beam structure under allometric cross-sectional growth. All data shown are generated in simulations using the rescaled beam model presented above. Force values are de-dimensionalized 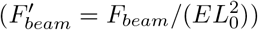 as in Fig. 5). For (a-c), the left column (sublabeled “1”) show the force scaling curves for various allometries in (a) *α*, (b) *β*, and (c) *s*. The right column (sublabeled “2”) show the same data normalized by the cross-sectional area scaling rate 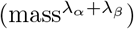, representing the residual force scaling associated with the correction term in Eqn. 33. Gray dashed lines indicate the scaling curves expected for a spring-like force scaling. (a) *λ*_*a*_ *∈* [0.25, 0.5], *λ*_*β*_ = *λ*_*s*_ = 1*/*3. (b) *λ*_*b*_ *∈* [0.25, 0.5], *λ*_*α*_ = *λ*_*s*_ = 1*/*3. (a) *λ*_*s*_ *∈* [0.25, 0.5], *λ*_*α*_ = *λ*_*β*_ = 1*/*3.

In contrast, when *λ*_*β*_ *∈* [0.25, 0.5] was allowed to vary (*λ*_*α*_ = *λ*_*s*_ = 1*/*3), the beam force does not scale like a spring structure (Fig. 6(b1)). When normalized, the curves no longer collapsed to the same curve (Fig. 6(b2)). Instead, the normalized curves also scaled with mass, with the slope scaling directly with the value of *λ*_*β*_. A similar trend is seen when the arc length is scaled (*λ*_*s*_ *∈* [0.25, 0.5], *λ*_*α*_ = *λ*_*β*_ = 1*/*3), but with the residual scaling slopes now inversely proportional to *λ*_*s*_. Thus, under allometric thickness or arc length scaling, we observe residual force scaling that cannot be explained by the scaling of the CSA, as predicted by the energy analysis.

### 3.4 Beam forces can scale faster than spring forces even without taking a larger fraction of the total system mass

So far, we have been lax about the coupling of the dimensional scaling rates *λ*_*s*_, *λ*_*α*_, and *λ*_*β*_. This results in situations where the beam’s mass ends up taking a different fraction of the system’s total mass as the system grows. The beam’s volume (*V*_*beam*_) is approximately equal to the integral of the CSA along the beam:

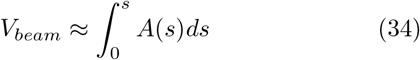

While not perfectly accounting for the effects of resting curvature, this is a useful approximation. Therefore, the beam’s volume scales as

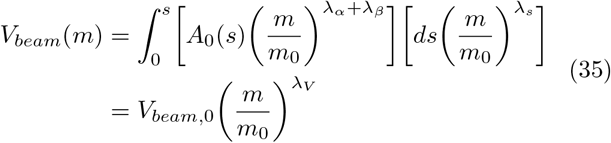

where *λ*_*V*_ = *λ*_*α*_ + *λ*_*β*_ + *λ*_*s*_ is the scaling rate of the beam’s volume, but *m* is the mass of the overall system. If the beam remains the same fraction of the system’s overall mass as the system grows with no changes in material density, then *λ*_*V*_ must equal 1, thus coupling the three axes as

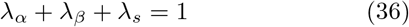

This can be enforced explicitly by requiring *λ*_*α*_ = 1 *− λ*_*β*_ *− λ*_*s*_. Applying this constraint to the scaling law for the beam force (Eqn. 33) and maintaining the assumption that *λ*_*x*_ = *λ*_*s*_, we obtain a constrained scaling law:

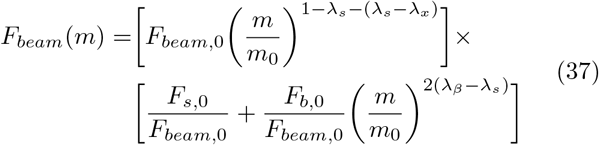

Under this more restricted scaling law, the leading term still scales with the CSA (*λ*_*A*_ = 1 *− λ*_*s*_), but we can see more clearly that the specific scaling of the cross-sectional dimensions does not impact this term. Instead, it can be fully determined by the scaling of the arc length and the isometric volume constraint. However, the correction term still depends explicitly on the scaling in the thickness direction. This also implies that, even for the case of 1) isometric arc length scaling (*λ*_*s*_ = 1*/*3), 2) isometric CSA scaling (*λ*_*A*_ = 2*/*3), and 3) no change in the beam’s relative mass (*λ*_*V*_ = 1), the beam forces can still scale differently than a corresponding spring-like structure. This could result from a differential scaling of the two cross-sectional directions. If the thickness were to scale at a faster rate than the width (*λ*_*α*_ = 2*/*3 *− λ*_*β*_, *λ*_*β*_ *>* 1*/*3) or more slowly (*λ*_*β*_ *<* 1*/*3), the force would correspondingly scale more quickly and more slowly, respectively. This occurs even in the case where standard spring scaling laws would predict the forces to scale as mass^2*/*3^ because the CSA grows isometrically.

### 3.5 Beam-like scaling changes the scaling of behavioral timing of quasistatic systems

The mechanics of beam-like structures allow their force properties to scale differently than a spring-like structure would, even under the same CSA scaling law. How does the allometric force scaling of beam structures impact the scaling of quasistatic behavior timing? How does this scaling differ from the scaling predicted for spring-like muscles? We investigated these questions using our simplified quasistatic agonist/antagonist model (See “Scaling of a quasistatic behavior“). For a 1-degree-of-freedom (1 DoF), quasistatic system actuated by an agonist/antagonist pair (Fig. 7(a1-a2)), steady-state cycle agonist activation can be written as a function of position (Eqn. 18, Fig. 7(b)). The duration of protraction (Eqn. 19) and retraction (Eqn. 20) can then be determined by the time constant of the muscle dynamics (*τ*) and the minimum and maximum positions of the mass (*q*_*a*_ and *q*_*b*_) during a steady-state cycle. To determine how these behavioral segment durations scale with mass, we must look at how time constants and activation scale with mass. If we assume time constants scale as a power law with parameter *λ*_*τ*_ (related to, e.g., the geometry of the motor neurons [24]), then Eqns. 19 and 20 can be rewritten as depending on animal mass as

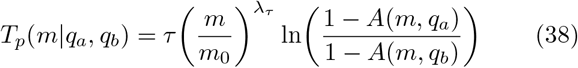

and

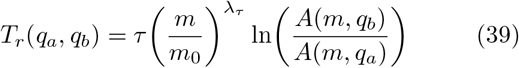

**Fig. 7:**
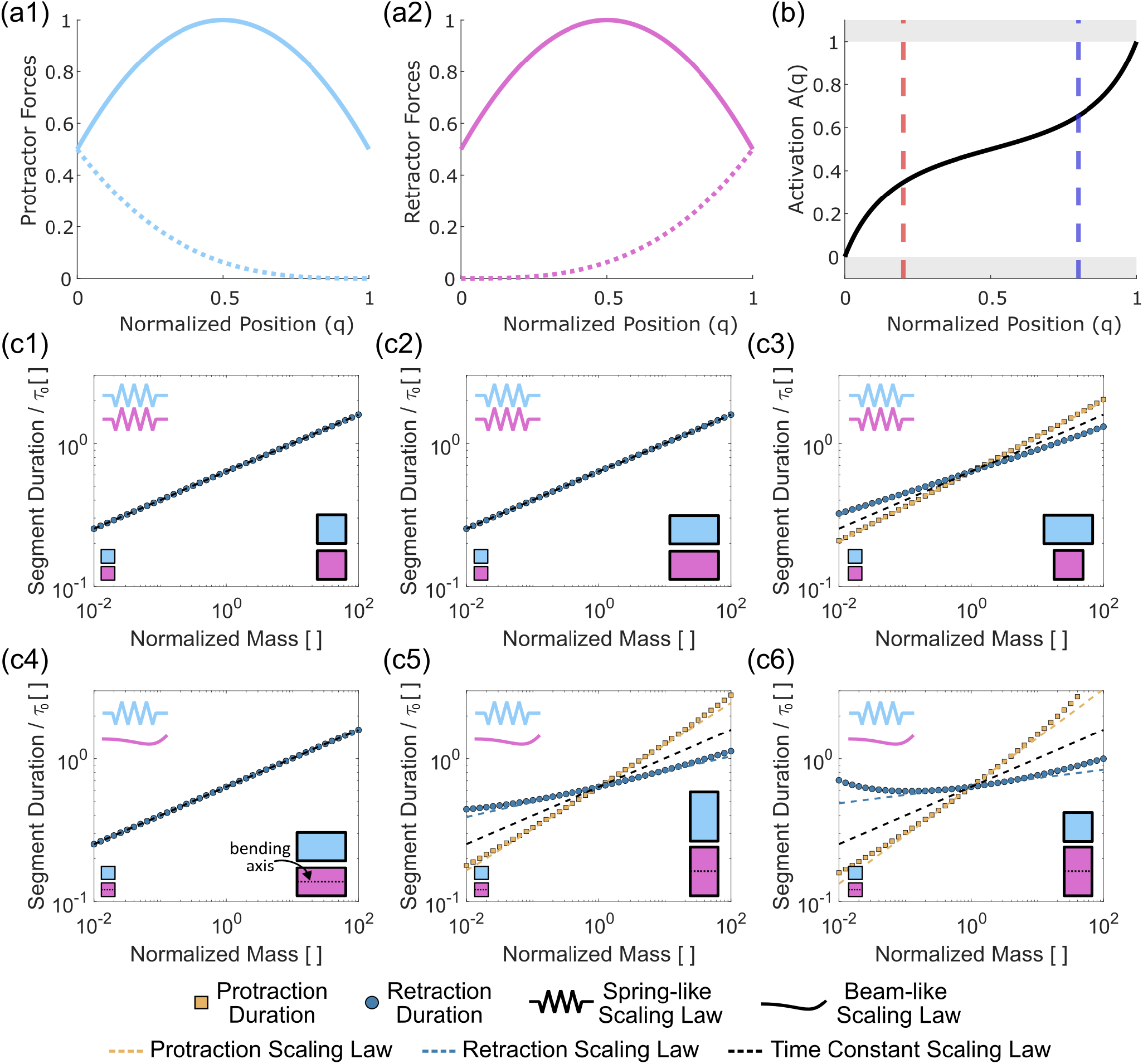
Behavioral scaling of a 1-degree-of-freedom quasistatic system. To demonstrate the qualitative predictions of our scaling analysis, we constructed a simplified version of the 1-degree-of-freedom model used during the analysis. Note: this does not attempt to replicate any particular biomechanical system. (a1-a2) Specific forms of the passive (dotted lines) and active (solid lines) force functions are implemented for the (a1) agonist and (a2) antagonist muscles. Here, 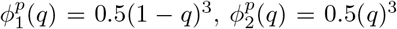 and 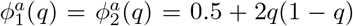. (b) Using these specific forms of the muscle force functions, we can calculate the activation-position function *A*(*q*) (Eqn. 18). The red vertical line shows the location of *q*_*a*_ = 0.2 and the blue line, *q*_*b*_ = 0.8. (c) Using the activation function from (b), we investigated the behavioral duration scaling for different prescribed scaling laws of muscle cross-sections and of the time constant *τ*. In (c), the cartoons at the bottom of the graphs schematically show the scaling of the structures’ cross-sections (*λ*_*α*_: horizontal; *λ*_*β*_: vertical). A square cross-section shows isometric growth, whereas a rectangle shows allometric growth. For each simulation, we set the time constant scaling parameter(*λ*_*τ*_ = 0.2) and the cross-sectional scaling parameters (*λ*_*α*_ and *λ*_*β*_) for each muscle. For (c1)-(c3), both muscles are assumed to be stretch-dominated and thus follow spring-like scaling laws for their force (*λ*_*j*_ = *λ*_*α*_ + *λ*_*β*_). For (c4)-(c6), the agonist muscle still behaves like a spring, while the antagonist muscle is replaced with a beam-like muscle (*λ*_2_ = *λ*_*α*_ + 3*λ*_*β*_ *−* 2*/*3). For the antagonist beam muscle, the thickness direction is scaled by *λ*_*β*_ and the width direction by *λ*_*α*_.

Activation is fully determined by the relative force contributions at a position *q*, so the scaling relationship for activation is determined by the scaling laws of the agonist/antagonist forces. If we assume the passive and active forces for a muscle scale identically and the agonist and antagonist scale as some power law with parameter *λ*_*j*_ for *j* ∈ [1, 2], then the mass-dependent activation equation can be written as

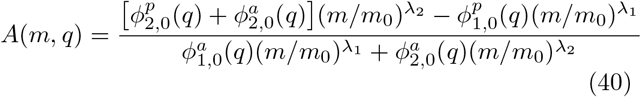

If both muscle forces scale at the same rate, *λ*_1_ = *λ*_2_, then the mass terms in the numerator and denominator factor and cancel, implying the activation is scale-invariant.

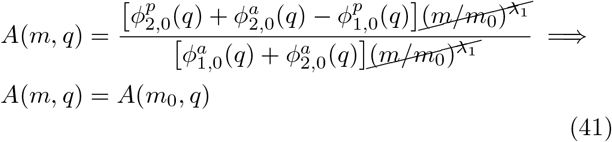

Therefore, if the agonist and antagonist forces scale identically, behavior duration scaling depends on the scaling of the muscles’ time constant (Fig. 7(c1)). Additionally, if the time constants of the muscle activation were also scale-invariant, this model would predict a scale-invariant behavioral duration for quasistatic systems. This scale-invariant activation and behavior duration has been previously derived for quasistatic systems [31].

This analysis did not require isometric growth, only that the muscle forces both grow at the same rate. This can be achieved by spring-like structures if the CSAs scale at the same rate, regardless of the scaling rates of the two cross-sectional axes (*λ*_*α*_ and *λ*_*β*_, Fig. 7(c2)). The only way for these behavioral durations to scale differently from the time constant in spring-like systems is for the two muscles’ cross-sections to scale differently (Fig. 7(c3)).

The same is not true when at least one of the muscles is beam-like. To illustrate this, we replaced the model’s antagonist with a muscle force that scales like a purely bending structure (*λ*_2_ = *λ*_*α*_ + 3*λ*_*β*_ *−* 2*/*3). For a system with a bending-dominated structure, having the CSAs of both muscles scale identically no longer guarantees the behavioral timings scale identically to the time constant. Instead, the directionality of the allometry now comes into play. If only the width scales allometrically (*λ*_*α*_*≠*1*/*3, *λ*_*β*_ = 1*/*3), then spring-like scaling still exists (Fig. 7(c4)). However, if thickness scales allometrically (*λ*_*β*_*≠*1*/*3), then, even for the same CSA scaling, the behavioral durations will no longer scale like the time constant (Fig. 7(c5)). Finally, when two systems (one with a spring-like retractor (Fig. 7(c3)) and one with a beam-like retractor (Fig. 7(c6))) share the same cross-sectional allometry, the beam-like system can demonstrate greater deviations from isometry (seen in the two plots as the differences in slope relative to the black dashed line). In the specific example shown here (Fig. 7(c6)), modest changes from isometry in geometric scaling rates can lead to a retraction scaling law that is nearly scale invariant (blue), while protraction duration (gold) scales far more quickly than the scaling of the activation dynamics (black). In general, because of the greater flexibility in their force response under different geometric scaling laws, beam-like structures allow for a wider range of possible behavior duration scaling laws than are allowed by purely spring-like muscles and elastic elements.

## 4 Discussion

### 4.1 The force scaling of bending biomechanical structures integrates mass, geometry, and loading conditions

The size and speed of animals and their behaviors are intrinsically connected to the governing physical forces with which their bodies interact [38, 7, 34, 8, 41, 23]. Studying how size, either across species in comparative studies or throughout growth within species, allows biologists and engineers to understand how and why different animals generate and control their behaviors. Animals must overcome the elastic and viscous forces of their constituent tissues and those imposed by the environment [6], which vary with size, requiring corresponding changes to the neural systems controlling behaviors [17]. For spring-like muscle and passive elastic forces, one only needs to consider how the CSA grows with mass [45, 15], regardless of the specific assumed or measured scaling law. Scaling relationships do not change if the geometry of the muscle or its loading conditions change as the body grows.

The same is not true for bending structures. While a beam’s stretching force still scales depending only on the CSA, the scaling of the bending force depends on the particular morphology of the cross section and the direction of bending because of the position-dependent term in the bending stiffness equation (*β*^2^ or *α*^2^, depending on the bending direction, Eqn. 10). If the morphology of a beam’s cross-section changes with mass, even for an isometric scaling of the cross-sectional area, the bending stiffness (and consequently the force) will scale differently because the distribution of the beam’s material has changed (see the Appendix for an example using a circular and annular beam). As discussed in the model derivation, the axial stiffness depends on the total amount of material, but the bending stiffness depends on the spread of the material distribution. Change that spread and the bending properties change as well. This effect cannot currently be captured in the force scaling law (Eqn. 33) as presented because of the assumed geometric scaling law. However, these morphology-based changes in the bending stiffness would lead to a similarly factorable scaling law, with the correction term depending on the specific morphology changes.

The bending force also depends on the arc length of the beam, even if the stroke length of the muscle applying the bending force and arc length scale identically. This is because the arc length of the beam acts as a moment arm for the muscle/tissue applying the bending force to the beam. If the moment arm scales faster than isometry, then a muscle will have to apply less force to the beam to cause the same level of bending.

Finally, the beam force scaling law (Eqn. 33) is not a simple power law, but a weighted power law. The weights come from the relative contributions of stretching and bending forces to the total force. These contributions will depend both on the overall geometry of the bending structure and on the loading condition. A beam loaded along its length will not bend (unless it buckles), but a beam loaded perpendicularly to its length will bend much more than it strains. Thus, if the loading conditions change with mass as a result of, e.g., the morphological remodeling of other anatomy, the resulting force scaling law will also change, even with no change to the anatomical scaling law. Similarly, if the same biological beam is loaded differently in two different behaviors, then the force scaling law of that beam may be different for each of those two behaviors.

In summary, when investigating the scaling of animals and behaviors that rely on bending structures, the focus cannot be solely on their anatomy but must be more holistic, incorporating changes in geometry and how structures interact with each other.

### 4.2 Allometric scaling of bending structures may shift behaviors, limit size, and provide material-efficient scaling of energy-transforming systems

If a bending structure exhibits an allometry in its thickness, the force it produces scales faster than the spring-like muscles and structures acting on it, even if both share the same cross-sectional allometry. Therefore, something must shift to compensate for this force imbalance. We demonstrated that muscle activation in a quasistatic system can scale with mass because of force scaling imbalances. However, this can only occur until the weaker muscle reaches its maximum activation (*A* = 1). At this point, the range of motion would begin to decrease or shift as the equilibrium position between agonist and antagonist elements changes. In our analysis of the 1-DoF translational model, we assumed the range of motion did not change as the mass changed. However, if a structure does demonstrate allometric growth, then this assumption may no longer be valid. Either the animal may shift its kinematics with size to maintain a similar neural control scheme, or it may be forced to shift when the weaker muscles reach their maximum activation. Alternatively, if a particular range of motion is critical to a behavior’s effectiveness, then an allometrically scaling bending structure may contribute to an animal’s maximum allowable size. That is, if the animal becomes too big, and consequently, its allometrically scaled bending structures become too stiff for its muscles to move through their range of motion, then this animal may no longer be able to perform its critical functions (e.g., feeding or locomotion). Thus, animals may need to balance the force gains provided by allometrically scaling beam thickness with the ability of their other muscles to deform those structures.

While allometric thickness scaling could limit an animal’s maximum size, it could also provide a space- and material-efficient method to scale the energetics of an elastic structure. The asymmetry with respect to the thickness direction in the bending stiffness causes the elastic energy stored by the beam (for a given deflection) to scale more quickly than a spring system under the same growth law. This means allometrically scaled beams will increase in their energy density relative to a spring of the same relative dimensions. Consequently, less material (and volume) would need to be added to the elastic structure for a required increase in maximum elastic energy storage. This could be relevant to latch-mediated spring-actuated (LaMSA) systems [25], like the jumping structures of grasshoppers [32] and planthoppers [4]. In these animals, the energetics of behaviors are dictated by the energy density of their passive structures and the muscles loading them. If the energy density of these LaMSA structures does not scale with mass, then we would expect behavioral parameters (e.g., take-off velocity, jump height) to be scale-invariant [39]. However, for some insects, takeoff velocities change with maturation in order to take flight as adults [21]. When observing the take-off velocities of juvenile nymphs and adult grasshoppers, the velocities were scale-invariant in the nymphs and increased when they reached their adult stage [10, 21]. This would require a morphological and scaling shift between nymphs and adults to allow for the energy density of the jumping mechanisms to increase. This would be achieved for the least amount of added mass by thickening (in the case of grasshoppers) the semilunar process, and, indeed, an allometric thickening of the cuticle is observed in adult grasshoppers relative to their nymph stages [11]. For LaMSA systems for which behavioral metrics scale with mass or developmental stage, it is important to investigate possible allometrically scaling bending structures.

### 4.3 Extensions of scaling analyses to plates and shells

This analysis of bending structures presented here was limited to beam-like structures, that is, those whose geometry can be approximated by a single primary topological dimension [2]. This may be an appropriate simplification for many biological structures (e.g., ribs of vertebrates, pleural arches of planthoppers, arms of cephalopods). Additionally, our own previous work has shown beam theory’s applicability to biological structures that, while not traditionally beam-shaped in their anatomy, deform by bending and stretching within a single plane [3]. However, many biomechanical structures that make use of bending are not topologically 1-dimensional. For example, many shells and exoskeletons are very thin relative to the dimensions of their surface area, thus allowing them to bend. However, rather than one topological dimension characterizing their geometry (i.e., arc length), the two topological dimensions of their surface area must be accounted for. Thus, these structures would be better approximated using mechanical plates or shells. Both plates and shells are topologically 2-dimensional mechanical structures that, like beams, can stretch and bend in 3D [1]. Their governing equations are of a similar form to those of beams, though with an additional dimension to consider. These structures were not investigated here because of their additional computational complexity. However, the bending and axial stiffness parameters along each perpendicular direction of these structures, and the expression for bending energy, present similar asymmetries with respect to thickness and arc length to those of beams. Therefore, while the specific scaling relationships for beams and plates may differ, many of the qualitative predictions that were made for beams (e.g., non-spring-like scaling only under allometric thickness growth) will still apply to plates. The additional complexities to the scaling relationships of plates and shells warrant further investigation, particularly as these structures are vital to many aspects of animal behavior.

### 4.4 Limitations and Future Directions

The current analysis produces some key qualitative predictions about the force scaling of bending-dominated structures. However, some limitations exist in the current approach. First, the analysis assumes a linear constitutive model for the biological materials. While this is appropriate for some biological tissues, like bone and cuticle, most (if not all) soft biological materials demonstrate meaningful nonlinearities in their constitutive relationships [18, 16]. Whether or not these nonlinearities affect the resulting force properties of the structures depends on the magnitudes of the structure’s deformations, but often these nonlinearities can be behaviorally important [14, 40]. Future investigations should more directly address these nonlinearities and the possible tensile/compressive modulus asymmetry. Additionally, here we only consider the axial and bending stiffness terms. However, for beams (and plates) with heterogeneous stiffness distributions, strain-bending coupling terms emerge [2]. Corresponding “coupling” force terms will also emerge in the force scaling weighted power law. These coupling parameters are position-dependent (like the bending stiffness) and, therefore, will introduce additional non-spring-like scaling. Thus, this analysis may be considered a minimal example of the possible non-spring-like scaling.

Finally, the assumed loading conditions place the loading force external to the bending structure (i.e., a muscle pulls on a distinct bending structure). However, these beam models apply to, and have been used extensively to model, the mechanics of muscular hydrostats [5, 43, 19]. These beam-like hydrostats (e.g., octopus arms and elephant trunks) are not always loaded by external structures but by intrinsic muscles throughout the structure. These muscles apply axial forces, bending/twisting moments, and stiffen the structure transiently and regionally [22]. Knowledge of how stiffness and muscle forces vary spatially is required to investigate the impacts of these muscles on scaling. Given the importance of octopus arms and elephant trunks as models of unconfined hydrostats [3] and as inspiration for soft robots, this would be a worthwhile investigation. This analysis also focused only on the effects of stretching and bending on the force output of these structures. For soft beams, additional deformations (like twisting, shearing) are behaviorally relevant, and thus more detailed beam theories (such as Cosserat theory [12]) should be analyzed within this framework.

## 5 List of symbols and abbreviations

*s*: arc length position along a structure
(*α, β*): cross-section coordinates
*N*(*s*): passive axial force generated by a beam or spring
*M*(*s*): passive bending moment generated by a beam
*µ*(*s*): axial stiffness
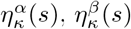: bending stiffness about the *α, β* axis
*ϵ*(*s*): axial strain
*κ*(*s*): curvature
Ω(*s*): spatial region defining the cross section
*E*(*s, α, β*): Young’s modulus
*m*: characteristic mass of the system
*λ*_*j*_: scaling power law exponent for the variable *j*
*q*: normalized position of a 1-degree-of-freedom quasistatic system
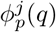: passive force from muscle *j* ∈ [1, 2] at *q*
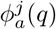: active force from muscle *j* ∈ [1, 2] at *q*
*A*_*j*_(*t*): normalized muscle activation for muscle *j* [1, 2] at time *t*
*N*_*j*_(*t*): normalized neural input to muscle *j* ∈ [1, 2] at time *t*
*τ*: time constant of muscle activation dynamics
*T*_*p*_: duration of protraction
*T*_*r*_: duration of retraction
*T*_*c*_: duration of full behavioral cycle
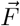: force vector applied to the tip of a biomechanical structure
*U*_*beam*_: total elastic energy stored in a beam
*U*_*b*_: elastic energy stored by changing the curvature of a beam
*U*_*s*_: elastic energy stored by changing the length of a beam
*F*_*beam*_: force at the tip of a beam generated by deforming the beam
*F*_*b*_: force at the tip of a beam associated with a change in curvature
*F*_*s*_: force at the tip of a beam associated with a change in length
*A*(*s*): cross sectional area
*V*_*beam*_: total volume of a beam structure

For all variables, the subscript “, 0” indicates values in the reference system.

## 6 Appendices

### 6.1 Application of the quasistatic analysis to a rotational system

Our quasistatic system analysis was based on a 1-degree-of-freedom (1DoF) translational system, where the mass translates along a single axis. The same analysis can be performed for a 1DoF rotational system (e.g., a knee joint, Fig. A1), and the same qualitative predictions are recovered. Here, we assume a link rotates around a fixed point, with angular position *θ* relative to the vertical axis. This link is acted upon by an agonist-antagonist pair, a flexor (“f”) and an extensor (“e”), both of which generate passive forces 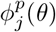 and active forces 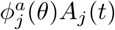. These are interpreted as they are in the translational case. The flexor acts at a distance *r*_*f*_ from the pivot point, and the extensor, at a distance *r*_*e*_. At each time point, the system must satisfy the following moment balance (analogous to the force balance of the translational system):

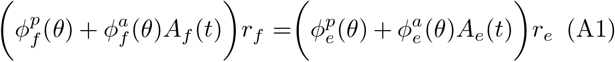

Assuming the same neural dynamics as in the original translational system (Eqn. 16) and that the two muscles act out of phase with each other (*A*_*e*_(*t*) = 1 *− A*_*f*_ (*t*) in steady-state cycles), we can solve for *A*_*f*_ as a function of angular position:

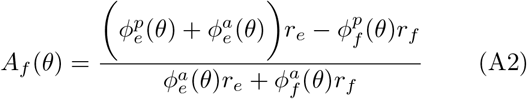

We can again cast the agonist activation as a function of position. The only differences between this equation and the equivalent equation for the translational system are the factors of *r*_*e*_ and *r*_*f*_, which are (to first approximation) fixed scaling coefficients that rescale the force terms but do not change the interpretation. From here, the rest of the analysis is identical, as are the qualitative predictions from the translational model.

**Fig. A1:**
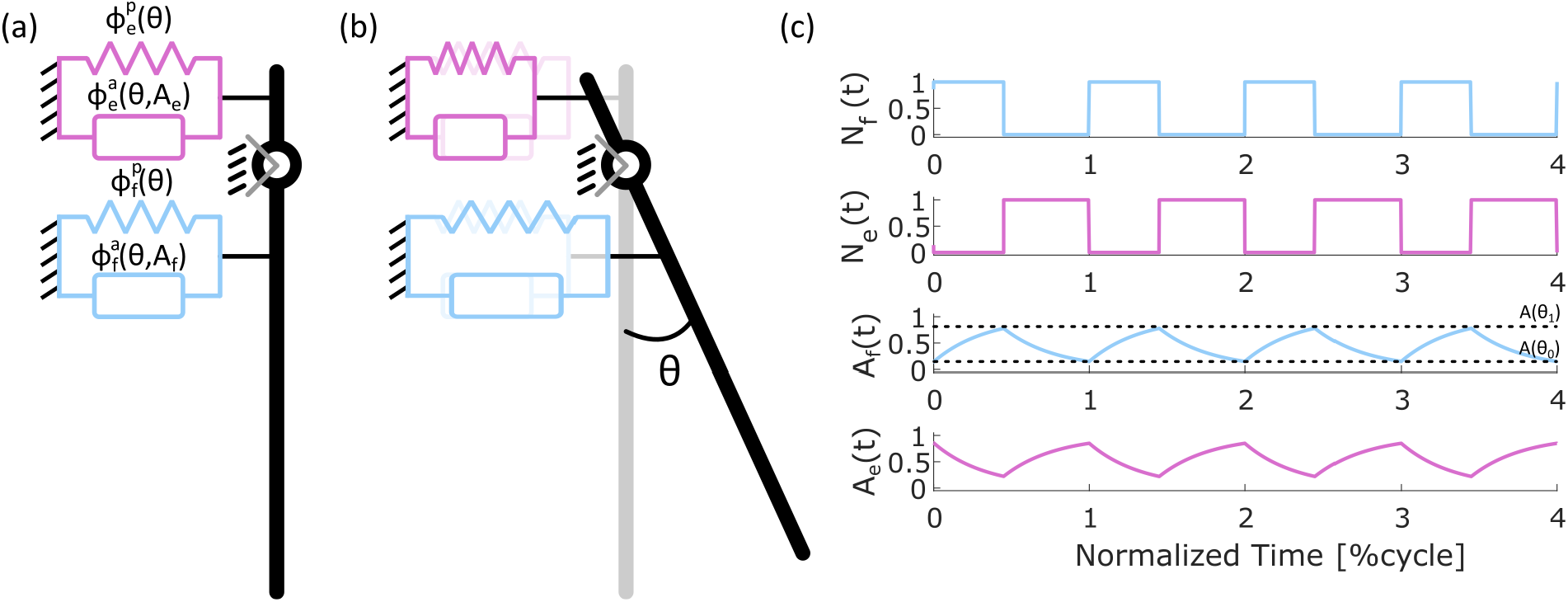
Quasistatic 1-degree-of-freedom rotational system. (a) As with the 1-DoF translational system analyzed in the main text, here a 1-DoF rotational system (e.g., a leg joint) is actuated by an agonist/antagonist pair. These are labeled “e” for extensor and “f” for flexor. These muscles can generate passive and active forces. (b) As the muscles activate, the leg swings through a range of angular positions (*θ*). (c) We again restrict the two muscles to receive neural stimulation (*N*_*j*_(*t*)) out of phase with each other such that their muscle activations (*A*_*j*_(*t*)) are also out of phase.

### 6.2 Impacts of Morphology Change on the Bending Stiffness

In the paper’s primary analysis, we investigate the impact of geometric scaling assuming no morphological changes or remodeling occur. Under this assumption, the cross-sections at all masses are identical except for the relative lengths of the two cross-sectional directions. However, as mentioned in the Discussion, the scaling of a bending structure can change under a morphology change, even if the CSA scaling law does not change. For example, suppose we have a structure for which, under a critical mass (*m*_*c*_), the cross-section is a circle. Then, above that critical mass, the cross-sectional morphology changes into an annulus. For example, earthworms and other animals with fluid at the center of their roughly circular bodies would be subject to differential scaling as a function of their growth. Another possible example from biology can be seen in leeches (*Hirudinea*), where small leeches are fairly round, whereas larger leeches are dorsoventrally flattened, which likely helps maintain flexibility around their primary bending axis (personal observations).

Let *R* be the radius of the circle under the critical mass, 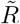 be the mean radius of the annulus, and Δ*R* be the half-width of the annulus (Fig. A2). The CSA of the circle is

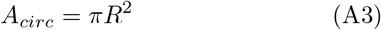

and for the annulus is

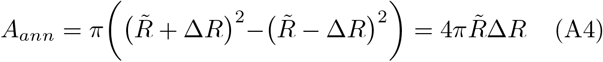

If we restrict Δ*R* to be a certain fraction of 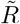 such that 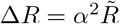, and if we require *A*_*circ*_ = *A*_*ann*_ for all mass ratios, then the dimensions of the annulus can be prescribed as:

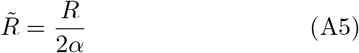

and

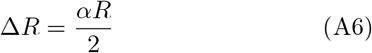

These equations hold true for any values of *R* and *α ≤*1. Next, we can calculate the axial and bending stiffness of the circular beam and the annular ring. For simplicity, we will assume the beams are made of a single uniformly distributed material with Young’s Modulus *E* such that *µ*_*ϵ*_ = *EA* and *η*_*κ*_ = *EI*, where *I* is the second moment of area. Because we require that the CSA of both beams be the same, the axial stiffness for both beams will also be the same. The second moment of area for a circle about its diameter is

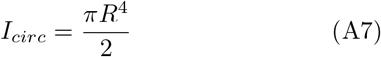

and for an annulus is

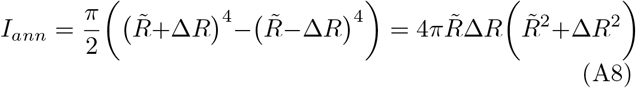

Substituting in the parameterized expressions for 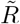 and Δ*R, I*_*ann*_ can be rewritten as

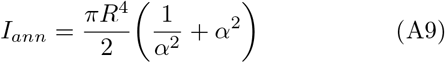

Even if we assume an isometric growth law, such that *R* mass^1*/*3^, then after the critical mass, when the beam’s morphology changes, the bending stiffnesses of the circular and annular beams will no longer be the same. If *α* is a fixed value, corresponding to a one-time remodeling event in the structure, this would only cause a shift in the bending stiffness. If instead the remodeling is continual, such that 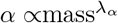, then there will not be a shift in the magnitude of the bending stiffness but in the scaling rate of the bending stiffness (Fig. A2). This is significant because now, even under CSA isometric growth, the bending stiffness (and therefore the resulting force) for the beam scales differently from a spring-like structure.

**Fig. A2:**
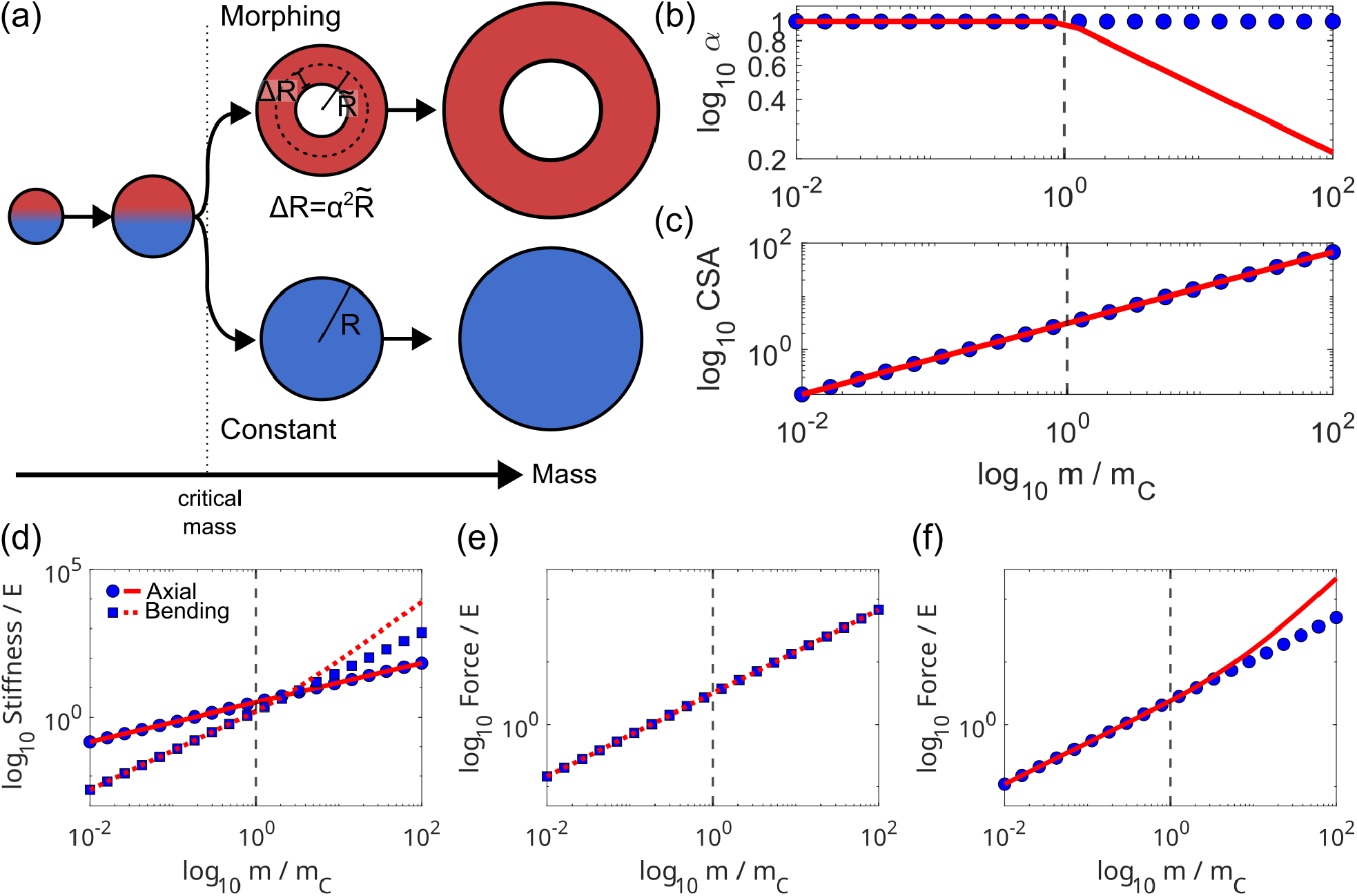
Impacts of morphology change on the force properties of bending structures. (a) Here, we analyze the effects of a beam’s cross-section morphing from a circle (blue) to an annulus (red). Note: CSA not to scale. (b) This morphology change is captured by a scaling of *α*. (c) Despite the morphology change, the CSAs of these beams are forced to scale (with mass) at the same rate. (d) While the axial stiffness scales identically between the morphing and constant beams, the bending stiffness scaling rate changes after the critical mass. (e) The spring-like forces produced by these structures scale identically despite the morphology change, but (f) the bending forces diverge above the critical mass.

## Acknowledgements

The authors would like to thank Dr. Roger Quinn and Dr. Ophelia Bolmin for their useful comments during the drafting of this paper.

## Competing Interestes

The authors have no competing interests to declare.

## Contributions

Conceptualization: MJB, SMR, GPS, HJC, VWW. Data Curation: MJB. Formal Analysis: MJB. Funding Acquisition: GPS, HJC, VWW. Methodology: MJB, VWW. Project Administration: MJB, VWW. Resources: VWW. Software: MJB. Supervision: HJC, VWW. Visualization: MJB. Writing – Original Draft: MJB, VWW. Writing – Review and Editing: MJB, SMR, GPS, HJC, VWW. 1

## Data availability

The code base developed for this article was based on the model code and numerical methods presented in Bennington et al. 2026 [3]. All code used to conduct the computational experiments was implemented in MATLAB r2024a. The code, including the necessary components of the code base from Bennington et al. 2026, is available for download on [GitHub](https://github.com/CMU-BORG/Scaling_of_BendingStructures). An archived version of the repository is also available through Zenodo (doi: 10.5281/zenodo.23249274). This article has no experimental data to share.

## Supplemental Materials

This article includes a Supplemental Materials document containing more detailed descriptions of the model’s governing equations and geometry, a primer on the multi-variable integrals used in calculating mechanical properties of these structures, and derivations/proofs for the equations utilized in the scaling analysis of the 1-DoF biomechanical system. 1

