## Supplemental Materials for "How bending scales: Geometric scaling differentially affects force responses in bending biomechanical structures"

### Supplemental Materials for
### Size, bending and behavior: Geometric scaling differentially affects force responses in bending biomechanical structures

Michael J. Bennington[1], Stephen M. Rogers[6], Gregory P. Sutton[7], Hillel J. Chiel[8,9,10], and Victoria A. Webster-Wood[1,2,3,4,5]

[1]Departments of Mechanical Engineering,
[2]Biomedical Engineering,
[3]Material Science and Engineering,
[4]McGowan Institute for Regenerative Medicine, and
[5]Robotics Institute, Carnegie Mellon University, Pittsburgh, PA, USA
[6]Department of Biology, University of Oxford, Oxford, UK
[7]School of Natural Sciences, University of Lincoln, Lincoln, UK
[8]Departments of Biology,
[9]Neurosciences,
[10]Biomedical Engineering, Case Western Reserve University, Cleveland, OH, USA

Submitted 17 Oct 2026

#### S1 Primer on multi-variable integrals

For the general parameter scaling law (Box 1), we make use of multi-variable integrals (that is, integrals taken over multiple spatial dimensions) to determine how material properties and geometry interact to produce the mechanical properties that we observe in biomechanical structures. When integrals are introduced, we typically talk about finding the area under a curve. However, a useful way to think about integrals is adding up the contributions of some property that is distributed over space. For example, the total mass ($M$) can be calculated by adding up the mass of smaller portions of the object ($M = \sum_i m_i$, where $m_i$ denotes the mass of the small portion we are considering). If we have a wire (Fig. S1(a)), we can define the linear density of the material (how much mass there is per unit length, $\rho_L(x)$) along the length of the wire. If we cut the wire into small sections, the mass of each segment is the linear density times the length of the small segment ($m_i = \rho_L(x_i)\Delta x$), and the total mass is

$$M = \sum_{i=1}^{N} \rho_L(x_i)\Delta x \tag{S1}$$

Here, $N = L/\Delta x$ is the number of segments into which we cut the wire, and $L$ is the length of the wire. If linear density changes over the length of the wire, then we can get a better approximation of the total mass by adding up smaller and smaller segments of the wire. We can then see that in the limit of the side length going to 0, we recover the definition of the 1-dimensional definite integral:

$$M = \lim_{\Delta x \to 0} \sum_{i=1}^{L/\Delta x} \rho_L(x_i)\Delta x \equiv \int_0^L \rho(x)dx \tag{S2}$$

This is a useful view of integration because it very naturally extends to multiple dimensions. For example, if we now consider a rectangular plate (Fig. S1(b)), we can define the area density (how much mass is there per unit of area, $\rho_A(x, y)$). We can then cut the plate into small squares of side length $\Delta x$ and $\Delta y$, and find the total mass

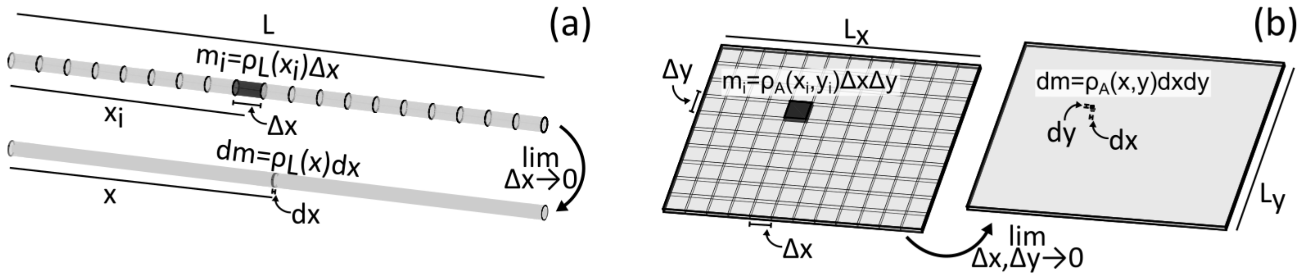


**Fig. S1**: **Single and multi-variable integration**. (a) 1-dimensional and (b) multi-dimensional integration can be envisioned as summing up small contributions of a distributed density function ($\rho_L$: linear density, $\rho_A$: area density). These multi-variable integrals can be used to calculate the beam mechanical parameters from the geometry of the beam's cross section and the distribution of Young's modulus. $m_i$: mass of segment $i$; $x_i$, $y_i$: position of segment $i$; $\Delta x$, $\Delta y$: side length of segment $i$. $dm$, $dx$, $dy$: infinitesimal mass and side length.

by first adding up the mass of each square in a row along the $x$ axis, and then together the mass of each row along the $y$ axis. Mathematically, this can be written as the nested sum

$$M = \sum_{j=1}^{M}\left[\sum_{i=1}^{N}\rho_A(x_i, y_j)\Delta x\Delta y\right] \tag{S3}$$

Here, $M$ is the number of rows along the $y$ axis. Just like with the wire example, we can get a better estimate of the total mass by decreasing the side length of our squares and taking the limit as those side lengths approach 0.

$$\begin{aligned} M &= \lim_{\Delta x,\Delta y\to 0}\sum_{j=1}^{L_y/\Delta y}\left[\sum_{i=1}^{L_x/\Delta x}\rho_A(x_i, y_j)\Delta x\Delta y\right] \\ &\equiv \int_0^{L_y}\int_0^{L_x}\rho_A(x,y)dxdy \end{aligned} \tag{S4}$$

This is the definition of the 2-dimensional (or double) integral. The same process could then be followed to define the multi-variable integral for higher numbers of dimensions, but for this paper, we will only need the double integral. For nice rectangular domains (or something similarly regular, like a circle), we can cleanly define the limits of integration. However, for many real-world geometries, these limits are not as obvious or cleanly defined. Therefore, for more general geometries, we can call the region over which we are integrating $\Omega$, and we can write the double integral of a property ($f(x,y)$) as

$$F = \iint_{\Omega} f(x,y)dxdy \tag{S5}$$

This is the notation we use in Box 1 whenever we integrate over a cross-section of a bending structure, since the specific geometry does not affect the scaling laws we derive.

### S2 Governing equations of spring and beam structures

For the scaling analysis of bending structures performed here, we utilized a planar form of geometrically exact Euler-Bernoulli beam theory. This is the geometrically nonlinear extension of traditional Euler-Bernoulli theory but still utilizes linear material models. Under Euler-Bernoulli assumptions, small segments of a beam can deform by changing their axial length or by curving into circular arcs (described by their curvature), but there is no shear in the cross section of the beam. Here, we will summarize the governing equations of the mechanical beam model. This quasistatic formulation is based on the model presented in Bennington et al. 2026 [SR1] and the dynamical model presented by Voesenek et al. 2020 [SR3]. For consistency, we use the variable and naming conventions from [SR1].

The configuration of a planar beam (Fig. **S2**(a)) is described by the position of the centerline ($\vec{x} = [x(s), y(s)]^T$) and the tangent heading angle of the centerline ($\theta(s)$). Here, $s$ is the position along the arc length of the beam in

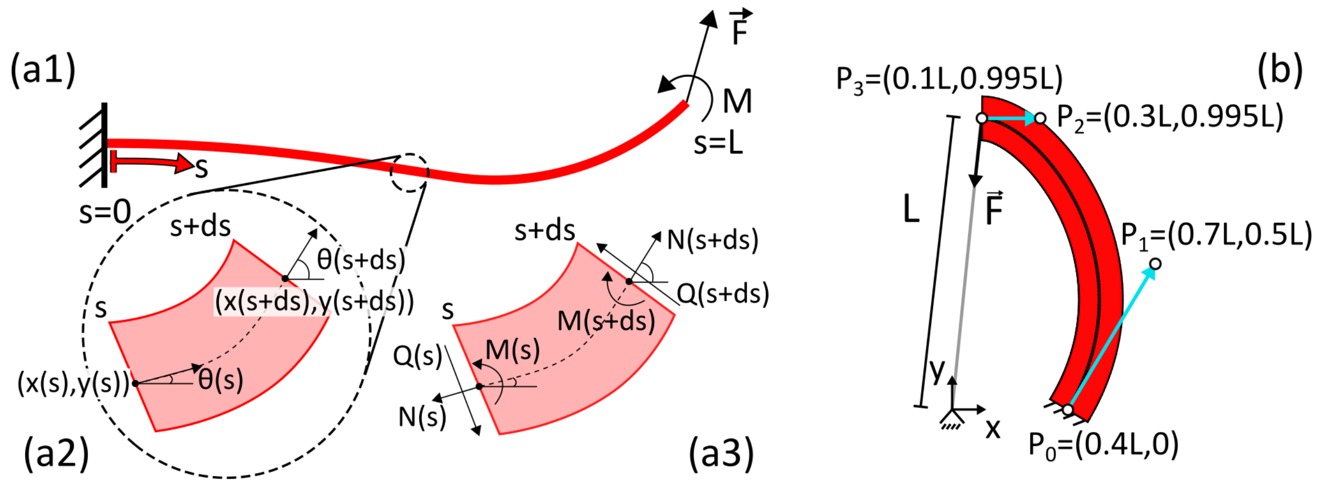


**Fig. S2**: **Mechanics of beam structures (a) and model geometry (b)**. (a1) A planar, geometrically exact Euler-Bernoulli beam is characterized by a 1-dimensional curve in the plane and parameterized along its arc length by the variable $s$. (a2) At each position, the configuration of the beam is captured in its planar position ($\vec{x} = [x(s), y(s)]^T$) and the tangent heading angle of the centerline ($\theta(s)$). (a3) To determine the equilibrium configuration of the beam subject to an external load, we must solve the force and moment balances at each position along the beam. $N(s)$: normal force, $Q(s)$: shear force, $M(s)$: bending moment. (b) The resting geometry of the beam used for the scaling analysis is defined by a Bézier spline. The four control points $P_0$-$P_3$ were set to approximate biological structures like the pleural arch of plant hoppers and the semilunar process of grasshoppers. The geometry is parameterized by the resting "muscle" length, $L$.

the resting configuration. When the beam is subjected to a set of boundary conditions (either kinematic boundary conditions constraining $x$, $y$, or $\theta$, or kinetic boundary conditions in the form of a force $\vec{F}$ or moment $M$), the equilibrium configuration of the beam is found by solving the simultaneous force and moment balance on each segment of the beam. For balances over infinitesimal segments of the beam (Fig. S2(a3)), this produces the coupled set of partial differential equations:

$$\frac{\partial}{\partial s}\Big(N(s)\hat{t}(s) + Q(s)\hat{n}(s)\Big) + \vec{f}(s) = \vec{0} \tag{S6}$$

and

$$\frac{\partial}{\partial s}M(s) + Q(s) + m(s) = 0 \tag{S7}$$

where $N(s)$, $Q(s)$, and $M(s)$ are the axial force, shear force, and bending moment distributions along the beam, $\hat{t}(s)$ and $\hat{n}(s)$ are the tangent and normal vectors of the beam, and $\vec{f}(s)$ and $m(s)$ are distributed body forces (e.g., gravity) and moments per unit length. For this analysis, we do not consider any distributed forces or moments, so these are equal to zero. These force/moment balance equations are coupled with a linear constitutive model that connects the deformations of the beam to its passive force and moment:

$$N(s) = \mu_\epsilon(s)\epsilon(s) \tag{S8}$$

and

$$M(s) = \eta_\kappa(s)\big(\kappa(s) - \kappa_0(s)\big) \tag{S9}$$

where $\mu_\epsilon$ and $\eta_\kappa$ are the axial and bending stiffness values whose scaling are examined in the main paper, and $\epsilon(s) = ||\partial\vec{x}/\partial s|| - 1$ is the axial strain and $\kappa(s) = \partial\theta/\partial s$ is the curvature. The shear force $Q(s)$ is determined from the force and moment balance. $\kappa_0(s)$ is the resting curvature of the beam under no loads. Finally, the heading angle $\theta(s)$ is constrained by the position of the beam through the equation

$$\theta(s) = \arctan\left(\frac{\partial y/\partial s}{\partial x/\partial s}\right) \tag{S10}$$

and the tangent and normal vectors are calculated as

$$\hat{t}(s) = \begin{bmatrix}\cos\theta(s)\\ \sin\theta(s)\end{bmatrix} \tag{S11}$$

and

$$\hat{n}(s) = \begin{bmatrix} -\sin\theta(s) \\ \cos\theta(s) \end{bmatrix} \tag{S12}$$

Eqns. S6-S10 are solved simultaneously subject to a set of six boundary conditions (three at each end of the beam) to determine the equilibrium position of the beam. Here, we discretize these equations using a finite difference approach and solve the resulting system of nonlinear equations using an adaptive form of Broyden's method [SR2] presented in Bennington et al. 2026 [SR1]. For details of the numerical methods, we refer the reader to that paper.

For the force-displacement experiments utilized in the scaling analyses, the $s = 0$ end of the beam (Fig. **S2**(b)) was fixed in place at its initial configuration ($\vec{x}(s = 0) = \vec{x}_0$; $\theta(s = 0) = \theta_0$), and a force was applied at the $s = L$ end ($[N(s)\hat{t}(s) + Q(s)\hat{n}(s)]_{s=L} = \vec{F}$; $M(s = L) = 0$). The force vector $\vec{F}$ points from the tip of the beam to the origin of the model (along the length of the "muscle"), and its magnitude was proportional to the length of the "muscle" relative to a desired set length:

$$\vec{F}(T) = -k\left(||\vec{x}(s = L)|| - (1 - 0.1T)L\right)\frac{\vec{x}(s = L)}{||\vec{x}(s = L)||} \tag{S13}$$

where $T \in [0, 1]$ is the normalized time of the 10% force ramp that is applied to the beam, and $L$ is the resting length of the "muscle." $k = (1 \times 10^3)\mu_\epsilon$ sets the stiffness of the constraint and incorporates the axial stiffness $\mu_\epsilon$ so that the same relative force is applied to the tip of the beam for a given value of $T$ across all beam sizes.

### S3 Beam model geometry and parameterization

The geometry of the beam model (Fig. **S2**(b)) used in the computational scaling analyses was defined by a 1-segment cubic Bézier spline, where the 2D points along the centerline of the beam ($\vec{x}$) are calculated as

$$\vec{x}(t) = \vec{P}_0(1 - t)^3 + 3\vec{P}_1(1 - t)^2t + 3\vec{P}_2(1 - t)t^2 + \vec{P}_3t^3 \tag{S14}$$

Here, $t$ is the parameter of the Bézier curve and does not correspond to the arclength parameter $s$ from the beam model. When generating the discretized geometry for the numerical calculations, the spline was sampled at each arc length interval (rather than equal intervals in $t$). The values of $\vec{P}_i$ were chosen to approximate the geometry of biological structures like the pleural arch or semilunar process. These values were parameterized by the rest length ($L$) of the "muscle" attached to the tip of the beam (light gray line). For beams of different mass ratios ($m/m_0$; see main document), $L$ was scaled isometrically to rescale the centerline geometry of the beam ($L(m) = L_0(m/m_0)^{1/3}$).

A single model geometry was used for all rescaled models by de-dimensionalizing the model. All lengths (including the width and thickness of the cross section) were normalized by $L$, and the Young's modulus was set to 1. However, this means that the simulation data for different masses effectively have different units. Therefore, to convert all of the simulation results (for force, axial stiffness, and bending stiffness) to the same normalized units, the results were rescaled by $(L/L_0)^2$, $(L/L_0)^2$, and $(L/L_0)^4$, respectively. Based on this rescaling, all force, axial stiffness values, and bending stiffness values have de-dimensionalized units of $EL_0^2$, $EL_0^2$, and $EL_0^4$, respectively.

### S4 Derivation of curvature scaling law

As part of the force scaling analysis, it is necessary to determine the scaling relationship between the tip deflection and the curvature of the beam. To approximate this, we can discretize the beam into $N$ segments, each with arc length $\Delta s = L/N$. The change in heading angle of each heading angle $\Delta\theta_i$ is related to the curvature and segment arc length as

$$\Delta\theta_i(s) = \kappa(s)\Delta s \tag{S15}$$

For equal arc length segments, the average heading angle change is then

$$\langle\Delta\theta_i\rangle = \langle\kappa\rangle\Delta s \tag{S16}$$

where

$$\langle\kappa(s)\rangle = \frac{1}{L}\int_0^L \kappa(s)ds \tag{S17}$$

is the average curvature along the beam. Next, the tip deflection of each segment, relative to the last segment, is equal to

$$|\Delta x_i| = \sin(\Delta\theta_i)\Delta s \tag{S18}$$

which, for small changes in heading angles reduces to

$$|\Delta x_i| \approx \Delta\theta_i \Delta s \rightarrow \langle|\Delta x_i|\rangle = \langle\Delta\theta_i\rangle\Delta s = \langle\kappa\rangle\Delta s^2 \tag{S19}$$

Therefore,

$$\langle\kappa\rangle = \frac{\langle|\Delta x_i|\rangle}{\Delta s^2} \tag{S20}$$

The total tip deflection is proportional to the segment tip deflections ($|\Delta x| \propto \langle|\Delta x_i|\rangle$), and the segment arc length is proportional to the total arc length ($\Delta s \propto L$) so

$$\langle\kappa\rangle \propto \frac{|\Delta x|}{L^2} \tag{S21}$$

Therefore, if $|\Delta x|$ scales as mass$^{\lambda_x}$ and $L$ scales as mass$^{\lambda_s}$, then

$$\boxed{\langle\kappa\rangle \propto \text{mass}^{\lambda_x - 2\lambda_s}} \tag{S22}$$

### S5 Proof for Equation 17

Let two signals $A(t)$ and $B(t)$ act as first order filters of inputs $N_A(t)$ and $N_B(t)$ such that

$$\frac{dA}{dt} = \frac{1}{\tau}\big(N_A(t) - A(t)\big) \tag{S23}$$

and

$$\frac{dB}{dt} = \frac{1}{\tau}\big(N_B(t) - B(t)\big) \tag{S24}$$

Let these two filters share a time constant $\tau$ and be activated out of phase with each other such that $N_B(t) = 1 - N_A(t)$. Therefore;

$$\frac{dB}{dt} = \frac{1}{\tau}\big(1 - N_A(t) - B(t)\big) \tag{S25}$$

If we consider the quantity $A(t) + B(t)$, its rate of change can be written as:

$$\frac{d}{dt}\big(A(t) + B(t)\big) = \frac{dA}{dt} + \frac{dB}{dt} \tag{S26}$$

Plugging in Eqns. S23 and S25 yields

$$\begin{aligned} \frac{d}{dt}\big(A(t) + B(t)\big) &= \frac{1}{\tau}\Big(N_A(t) + 1 - N_A(t) - \big(A(t) + B(t)\big)\Big) \implies \\ \frac{d}{dt}\big(A(t) + B(t)\big) &= \frac{1}{\tau}\Big(1 - \big(A(t) + B(t)\big)\Big) \end{aligned} \tag{S27}$$

Thus, regardless of the activation $N_A(t)$, the quantity $A(t) + B(t)$ will always approach 1 as $t \rightarrow \infty$. If $N_A(t)$ oscillates periodically, in steady-state cycles, $A(t) + B(t) = 1 \implies B(t) = 1 - A(t)$.

### S6 Derivation of Protraction and Retraction Duration

Let the filter $A(t)$ oscillate between two values $A_0$ and $A_1$ where $A_0 < A_1$. Let the filter $A(t)$ receive a normalized input $N(t) \in [0, 1]$ such that the governing equation of the filter can be written

$$\frac{dA}{dt} = \frac{1}{\tau}\big(N(t) - A(t)\big) \tag{S28}$$

If, during activation ($A_0 \rightarrow A_1$), the normalized input is maximal ($N(t) = 1$), the governing equation reduces to

$$\frac{dA}{dt} = \frac{1}{\tau}\big(1 - A(t)\big) \tag{S29}$$

where $A(t=0)=A_0$. This equation has the solution

$$A(t) = 1 - (1 - A_0)e^{-t/\tau} \tag{S30}$$

At the end of activation (at $t = T_p$ following the nomenclature from the main text), $A(T_p) = A_1$. Substituting this into Eqn. S30, we obtain

$$A_1 = 1 - (1 - A_0)e^{-T_p/\tau} \implies T_p = \ln\left(\frac{1 - A_0}{1 - A_1}\right) \tag{S31}$$

If, during relaxation ($A_1 \to A_0$) the normalized input is minimal ($N(t) = 0$), the governing equation reduces to

$$\frac{dA}{dt} = -\frac{1}{\tau}\big(A(t)\big) \tag{S32}$$

where $A(t=0)=A_1$. This equation has the solution

$$A(t) = A_1 e^{-t/\tau} \tag{S33}$$

At the end of relaxation (at $t = T_r$), $A(T_r) = A_0$. Substituting this into Eqn. S33, we obtain

$$A_0 = A_1 e^{-T_r/\tau} \implies T_r = \ln\left(\frac{A_1}{A_0}\right) \tag{S34}$$
